# Rethinking habitat occupancy modeling and the role of diel activity in an anthropogenic world

**DOI:** 10.1101/2021.06.30.450589

**Authors:** Kimberly Rivera, Mason Fidino, Zach J. Farris, Seth B. Magle, Asia Murphy, Brian D. Gerber

**Affiliations:** Department of Natural Resources Science, University of Rhode Island, Kingston, RI 02882, USA; Urban Wildlife Institute, Lincoln Park Zoo, Chicago IL, USA; Department of Health and Exercise Science, Appalachian State University, Boone, NC 28608, USA; Department of Environmental Studies, UC Santa Cruz, Santa Cruz, California, USA

**Keywords:** camera traps, diel, habitat, multi-state, occupancy, spatial-temporal, temporal activity

## Abstract

Current methods to model species habitat use through space and diel time are limited. Development of such models is critical when considering rapidly changing habitats where species are forced to adapt to anthropogenic change, often by shifting their diel activity across space. We use an occupancy modeling framework to develop a new model, the multi-state diel occupancy model (MSDOM), which can evaluate species diel activity against continuous response variables which may impact diel activity within and across seasons or years. We used two case studies, fosa in Madagascar and coyote in Chicago, USA, to conceptualize the application of this model and to quantify the impacts of human activity on species spatial use in diel time. We found support that both species varied their habitat use by diel states—in and across years, and by human disturbance. Our results exemplify the importance of understanding animal diel activity patterns and how human disturbance can lead to temporal habitat loss. The MSDOM will allow more focused attention in ecology and evolution studies on the importance of the short temporal scale of diel time in animal-habitat relationships and lead to improved habitat conservation and management.

## 1. INTRODUCTION

*“No description of where an animal lives and what it does can be complete without considering when the activity takes place.”* -- (Enright, 1970)

Understanding a species’ or communities’ habitat is one of the most fundamental aims of ecology (Mitchell, 2005) and conservation (Campomizzi et al., 2008). Historically, habitat was defined by Odum et al. (1971) as “the place where an organism lives, or place where one would go to find it.” This fundamental definition has evolved in recent years to address both space and time, such as “a description of a physical place, at a particular scale of space and time, where an organism either actually or potentially lives” (Kearney, 2006). Redefining habitat to encompass both spatial and temporal scales has allowed studies to improve hypotheses of how organisms interact with their environment (Kearney, 2006; Morano et al., 2019), which better recognizes how space and time are two fundamental axes of a species’ niche (Pianka, 1973).

Empirical knowledge of species’ habitat has grown with the development of spatial modeling, including species distribution (Segurado and Araújo, 2004), occupancy (MacKenzie et al., 2017), and resource selection models (Northrup et al., In Press). Inferences from these models have helped identify critical habitats of threatened species (Guisan et al., 2013), manage invasive species (Guisan et al., 2013), and understand how landscape structure (e.g., landcover) impact species habitat use (Angelieri et al., 2016; Hirzel et al., 2006). However, while the application of these models can identify fine scale spatial information of a species’ habitat, they focus on larger temporal patterns, such as seasonal or yearly scales (Fidino and Magle, 2017; MacKenzie et al., 2003). Species activity over diel time, typically described via defined modalities like diurnal or nocturnal (Anderson and Wiens, 2017), has a fundamental role in their space use (Pianka, 1973). These studies ignore this critical temporal period, making it difficult to understand how rapidly changing conditions and landscapes impact a species’ daily activity (Ellis et al., 2010; Gaston, 2019; Helm et al., 2017). The limited studies that do consider space use and diel activity, however, treat them separately and observations are not associated with a continuous response variable that can only provide descriptive inferences rather than an explicit estimation of hypothesized effects (James et al, 2013; Ridout and Linkie, 2009). For instance, when spatial inferences are made via occupancy modeling (Long et al., 2011) and diel temporal inference via circular kernel density methods (Ridout and Linkie, 2009). Therefore, these studies have focused on ‘average daily conditions rather than those prevailing at the time of day when individuals would tend to be most active’ (Gaston, 2019).

Evaluating space use in diel time is especially urgent given increasing anthropogenic pressures across landscapes globally (Ellis et al., 2010). If species can adjust their diel activity, then it and could be a mechanism by which they adapt to changing landscapes, climate, or ecological communities. For instance, meso- and large-carnivores have been found to increase their nocturnal activity in urbanized areas (Carter et al., 2012; Gehrt, 2007), likely to avoid when humans are most active (Gaynor et al., 2018). During hunting seasons, harvested species such as deer can become more nocturnal to avoid hunters (Kilgo et al., 1998). Animals may also change their diel activity in the presence of introduced species, as is the case with many mammals (ungulates, carnivores, and small mammals) who temporally avoid domestic dogs (Farris et al. 2015; Lenth et al. 2008). By modifying behavior across the 24-hour light-dark cycle, species can access space that would otherwise be inaccessible. This flexibility, however, may have physiological, morphological, or even ecological constraints, such as limited diel periods in which food is available for hunting or foraging (Kronfeld-Schor et al., 2017). Understanding a species’ spatial activity across diel-time use can therefore provide insight into these constraints, leading to a more complete understanding of where species live and how pressures impact their habitat. For example, a species may lose spatial resources altogether or lose spatial resources during a specific diel time period, such as hours when humans are most active (Ellington et al., 2020). Pumas, for instance, had diminished daily access to food resources in response to simulated human disturbance via playback (Smith et al., 2017). By considering spatial and temporal habitat jointly in a single modeling framework, we can explicitly evaluate hypotheses regarding how an animal’s relationship with the landscape changes as humans alter resources and the risk of obtaining those resources.

With increasing availability of camera traps, which allow for passive and continuous sampling of wild animal populations (Rovero et al., 2013), we also have increasing access to fine scale spatial-temporal data required for joint analyses of space use and diel activity. To advance theories of ecology and their application, we require a model which bridges gaps in existing capabilities by incorporating continuous covariates on diel behavior within a framework that accounts for variation in detectability across space and diel time. As such, our objectives were to, 1) develop static and dynamic occupancy models that incorporate diel activity information (multi-state diel occupancy models; MSDOM) and 2) apply these models to investigate how anthropogenic development and activity may simultaneously alter where and when species occur. We do so by presenting a case study on Madagascar’s largest endemic carnivore, the fosa (*Cryptoprocta ferox*), to demonstrate the static MSDOM, and a case study on the urban ecology of coyote (*Canis latrans*) to demonstrate the dynamic MSDOM. With our MSDOMs, and the growing availability of spatial and temporal data, it is possible to evaluate hypotheses on wildlife diel activity across space and through time, making it a major advancement over current methods (Azzou et al., 2021; Distiller et al., 2020).

## 2. MATERIALS AND METHODS

### 2.2 Multi-state diel occupancy models

#### 2.2.1 Static Model: a single season occupancy analysis

The MSDOM is a form of the multi-state occupancy model with state uncertainty (MacKenzie et al., 2009; Nichols et al., 2007) and is defined below with four states equivalent to the original co-occurrence model (MacKenzie et al., 2004) with two-species; the static model can also be understood as a special case of the species co-occurrence model by Rota et al. (2016) and the dynamic model a special case of Fidino et al. (2019). However, the MSDOM considers biologically important diel time periods for state segregation; this segregation can be based on any set of time periods of interest. In our case, sites are defined in one of four (*M* = 4) mutually exclusive states: 1) ‘no use’, 2) ‘day use’, 3) ‘night use’, and 4) ‘day & night use’. While these are coarse categorizations for diel behavior, these states provide us the ability to quantify the strength of drivers to diel shifts across continuous space and therefore identify biologically informed thresholds for species diel habitat use. Surveys are conducted over spatial locations, or camera trap sites (*i* = 1, *N*), which are independently sampled on *j* = 1,.., *K* occasions (e.g., days or weeks). Our state definitions do not follow a hierarchical ordering as commonly applied in multi-state occupancy models (Nichols et al., 2007) and implemented in R packages (unmarked; Fiske and Chandler, 2011). For example, if site *i* was observed in state 2, it precludes the site from ever being in state 3 as these states do not co-occur over a given survey period.

##### 2.2.1.1 Full Model (no covariates)

Let *ψ*^m^ be the probability that a site is in occupancy state *m* where **ψ** = [*ψ*^1^ *ψ*^2^ *ψ*^3^ *ψ*^4^] is the state probability vector, *ψ*^1^ = 1 – *ψ*^2^ – *ψ*^3^ – *ψ*^4^, and **1 o ψ** = 1 (see parameter descriptions in Appendix S1). The marginal occupancy probability (regardless of state) is *ψ*^•^ = *ψ*^2^ + *ψ*^3^ + *ψ*^4^. Then, let, 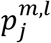 be the probability of observing the occupancy state *l*, given the true state is *m* in *j* survey. The detection probability matrix for survey *j* (assuming no site or survey variation) is *M* × *M* with the observed (columns) and true states (row) with rows summing to 1,

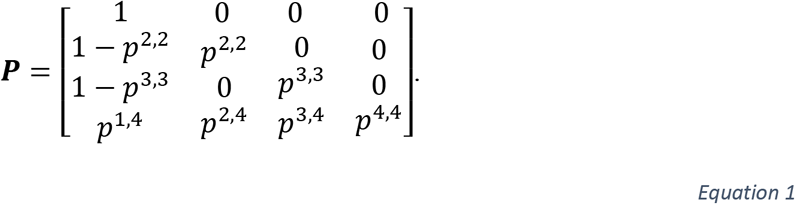

Together, the true occupancy state for site *i* is defined by the latent variable, ***z**_i_*~ Categorical(***ψ***) and the observed state in survey *j* is defined as, *y_ij_*~ Categorical (***P**_z_i__*). Taking a Bayesian modeling framework, we can assume diffuse prior distributions for model parameters as ***ψ, P***_4_ ~ Dirichlet (1,1,1,1) and *p*^2,2^, *p*^3,3^~ Beta (1,1). Note that in this full model, there is no relationship among state-specific detection probabilities (i.e., *p*^2,2^, *p*^3,3^, ***P***_4_) and occupancy probabilities (i.e., *ψ*^2^, *ψ*^3^, *ψ*^4^) across associated *M* states. Specifically, state 4 (‘day & night use’) occupancy and detection is not defined by state 2 (‘day use’) and 3 (‘night use’). This suggests that there is a fundamental difference between sites or species activity that occupy state 4. Species present during the ‘day & night’ state may be cathemeral, indicating they have intermediate adaptations allowing them behavioral flexibility to manage disturbance (Bennie et al., 2014). We can also estimate a species temporal use on the landscape by conditioning on species presence to examine how species navigate anthropogenic features via time partitioning. We do this by investigating an occupied state of interest over the sum of all occupied states. For example, the likelihood a species will use the ‘night’ state given it is present, is 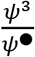.

##### 2.2.1.2 Reduced Model (no covariates)

The reduced model is a simpler parameterization that defines the occupancy and detection probabilities of state 4 (‘day & night use’) as the product of states 2 and 3. Therefore, we assume the diel time periods of ‘day & night’ are independent random events, allowing their probability products (detection and occupancy) to result in the probability of occurring or being detected during the ‘day & night’. Here, we can redefine our model in terms of the probability of using a site during the day, regardless of use at night (marginal probability; *ψ*^Day.M^) and the probability of using a site at night, regardless of use during the day (*ψ*^Night.M^). Our state occupancy probabilities are then,

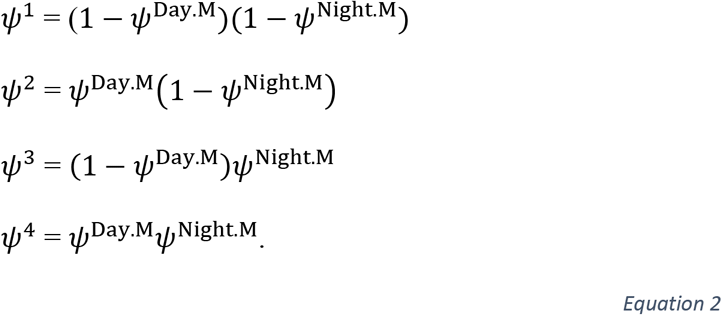

Similarly, we can define ***P*** using the probability of detection during the day (*p*^Day.M^) and night (*p*^Night.M^) as,

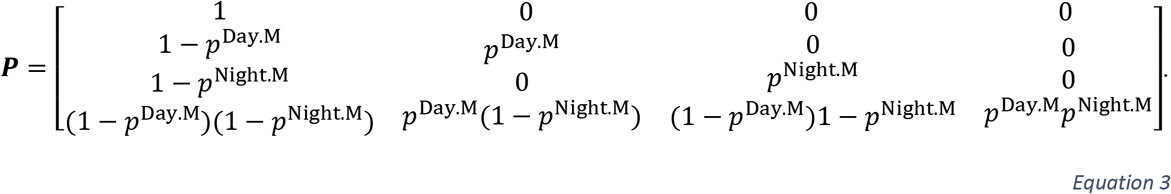

We can assume diffuse prior distributions for our reduced model parameters:

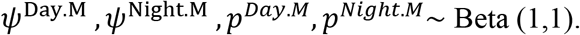

##### 2.2.1.3 Null Model

It is important to compare more complex models with one that does not consider diel time partitioning. This null model would thus be a single season occupancy model (MacKenzie et al., 2002), cast in a multi-state framework for model comparison purposes. Our state occupancy probabilities are then,

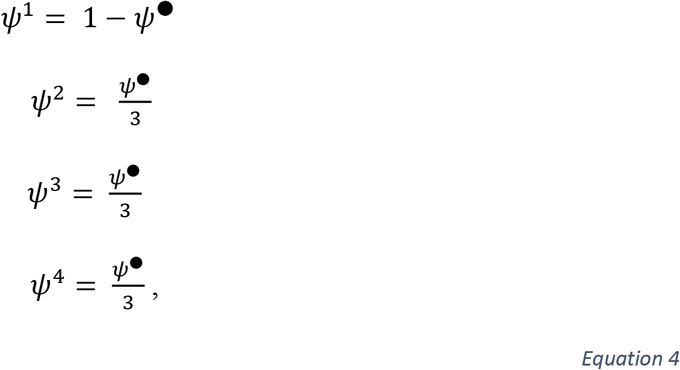

with the following detection matrix,

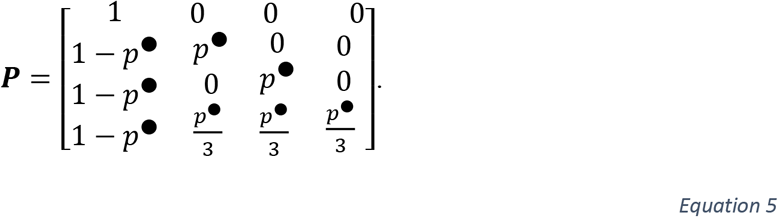

We can assume the following diffuse prior distributions for the null model parameters:

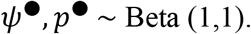

##### 2.2.1.4 Models with Covariates

All versions of the MSDOM (full, reduced, null) allow for the incorporation of site level covariates as explanatory variables of ***ψ*** and ***P*** and survey level covariates for ***P***. We use separate design matrices for modeling each state 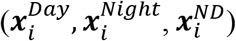 that for each site *i*, are 1 × *Q_m_* (the number of columns) and associated vectors of coefficients (**α**^*Day*^, **α**^*Night*^, **α**^*ND*^) that are *Q_m_* × 1. We link state-specific linear models with occupancy probabilities using the multinomial logit link. The full model with covariates is specified as,

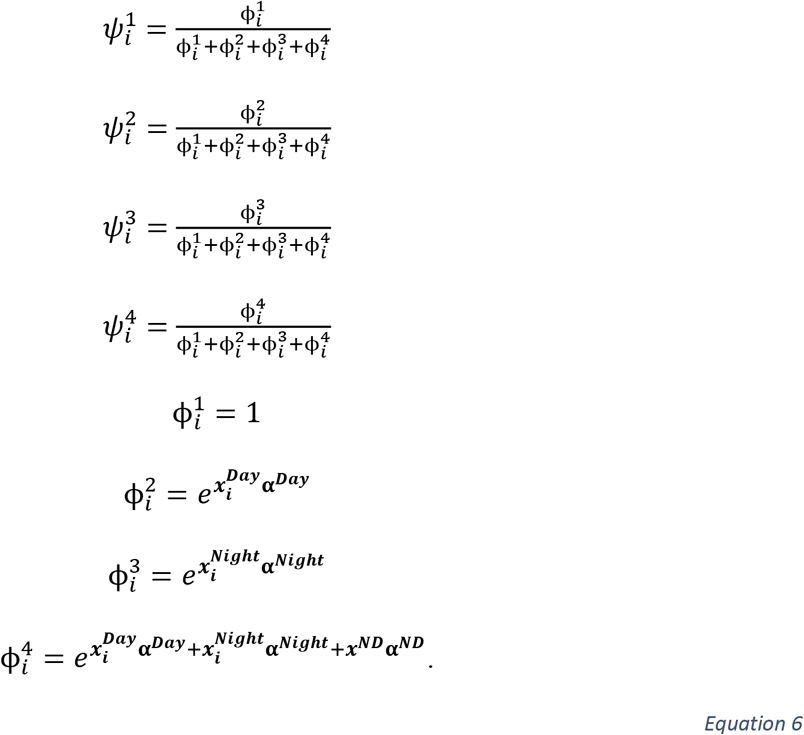

Here, 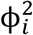 and 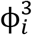 only contain first-order parameters, which respectively represent the log-odds a species occupies site *i* in either state 2 or 3 (i.e., they are associated to a single state). The parameter 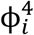, however, also contains second-order parameters 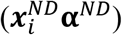, which represents the log-odds difference a species occupies site *i* in state ‘day & night’ relative to the aforementioned first-order parameters (see Dai et al., 2013). Thus, the second-order parameters for the ‘day & night’ state allows us to evaluate if this state is different than the day and night states combined. To specify the reduced model, we remove 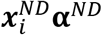 from the linear model on 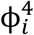. The null model with covariates is recast to leverage the unoccupied state equally to the combination of the identical, but multiple occupied states as,

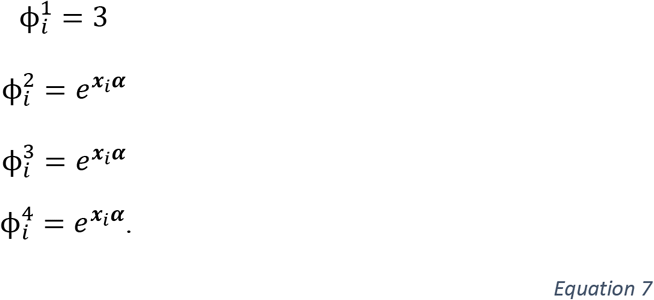

We can assume diffuse prior distributions for all coefficients as α_m_ ~ Logistic (0,1; Northrup and Geber, 2018). Including covariates on the detection matrix similarly uses the multinomial logit link (see Appendix S2).

#### 2.2.2 Dynamic Model: across season occupancy analysis

The dynamic MSDOM considers how site use at the diel scale changes over longer-time scales, such as seasons or years. The sampling protocol is identical to a static MSDOM except sites are sampled over *t* = 1,…, *T* primary sampling periods. Furthermore, we assume occupancy state, **z**_*i,t*_, depends on the state in the previous primary period, **z**_*j,t*−1_, which allows transitions to be estimated in terms of state-specific local colonization (*γ*) and extinction (ε) for all sampling periods save for the first. Instead, we estimate initial occupancy for the first sampling period like the static MSDOM. For all dynamic MSDOM, let ***τ*** be an *M* × *M* transition matrix whose rows sum to 1 and contains the rates that describe the probability a site either stays in the same occupancy state or transitions to a new state from one primary sampling period to the next.

##### 2.2.2.1 Full model (no covariates)

While the most general full model would independently estimate all *M* × *M* transitions among states, such a model may be difficult to fit with typical sample sizes from real world data. Thus, we imposed a few biologically reasonable constraints to reduce the number of model parameters and allow for more sparse, but realistic datasets to be used. For the full model, let ***τ*** be

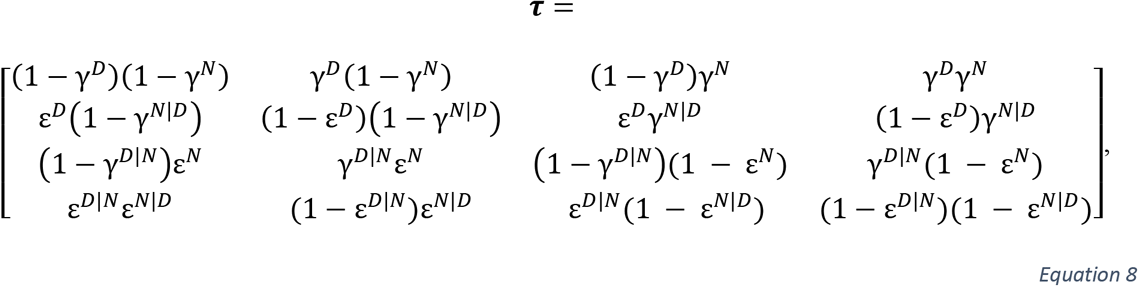

where the rows respectively describe state transitions from the four occupancy states. For example, the probability a site changes from state 2 (‘‘day use’) to 3 (‘night use’) is *τ*^2,3^ = ε^*D*^γ^*N*|*D*^, where ε^*D*^ is the site extinction probability in the ‘day use’ state and γ^*N*|*D*^ is the probability of colonization of the ‘night use’ state given ‘day use’ in the previous primary period. We assume that transitions depend on the state in the previous primary period, and that transitions from occupied states (i.e., 2, 3, or 4) may not be equivalent to transitions from the unoccupied state (i.e., state 1).

As with the full static MSDOM, the initial occupancy probability of the four states at *t* = 1 is 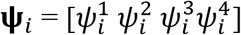. The latent state of the model is then **z**_*i*,1_ ~ Categorical (**ψ**_*i*_) for *t* = 1 and **z**_*i,t*_ ~ Categorical (*τ*_**z**_*i, t*−1__) for *t* > 1, where ***z***_*i,t*−1_ indexes the appropriate row of ***τ***. The observed state is specified like the full static MSDOM, except we index the observed data and latent state through time such that ***y**_ijt_* ~ Categorical (***P**_z_i,t__*), where ***P*** is Eq. 1 and ***z**_i,t_* indexes the appropriate row of ***P***. Finally, we assume the same diffuse prior distributions as the full static MSDOM for ***ψ*** and ***P*** while all colonization (***γ***) and extinction (**ε**) parameters have their own respective Beta (1,1) distributions.

##### 2.2.2.2 Reduced model (no covariates)

The reduced dynamic model is similar to the full dynamic model except initial occupancy becomes Eq. 2, ***τ*** lacks conditional parameters, and ***P*** becomes Eq. 3. Therefore, ***τ*** simplifies to

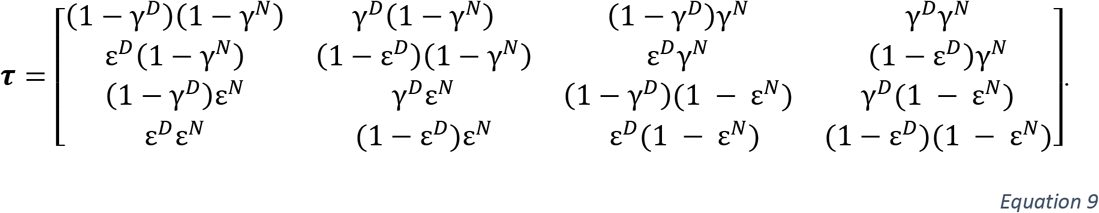

With the exclusion of conditional parameters, this model assumes that transitions between day and night are independent random events.

##### 2.2.2.3 Null model (No covariates)

Casting the dynamic MSDOM as a standard multi-season occupancy model requires splitting the associated colonization and extinction probabilities across each respective row of ***τ*** to ensure each row still sums to 1 such that,

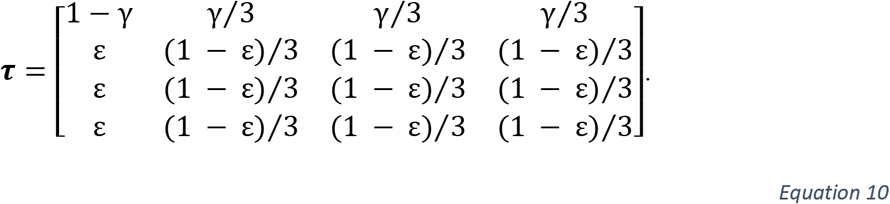

As with the static null MSDOM, initial occupancy becomes Eq. 4 and ***P*** becomes Eq. 5.

##### 2.2.2.4 Models with covariates

As with the static MSDOM, transition probabilities for each dynamic model can be made a function of covariates. To do so, we use separate design matrices for each model parameter which are 1 × *Q_m_* (e.g., 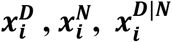, and 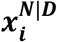) and associated vectors of coefficients that are *Q_m_* × 1 (e.g., ***b^D^, b^N^, d^D^, d^N^, g^D|N^, g^N|D^, h^D|N^***, and ***h^N|D^***). Temporal or spatiotemporal covariates may also be included in dynamic MSDOM, resulting in *T* × *Q_m_* design matrices for colonization, extinction, or detection parameters. Following Fidino et al. (2019), the linear predictors for the parameters of the full model are,

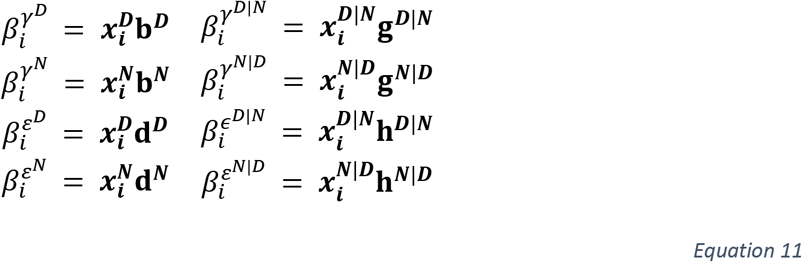

for the dynamic model, 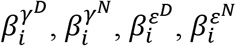 are first-order parameters while 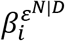, and 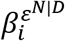 are second-order parameters. In this case, the second-order parameters are the logodds difference, given the presence of another state in either the current time step (*t*) for occupancy and detection or in the previous time step (*t-1*) for colonization and extinction. Let ***ω*** be a matrix with the same dimensions as ***τ*** that contains the linear predictors of the dynamic model. We set the diagonal of the matrix as the reference category so that transitions are estimated relative to a site staying in the same state from one time step to the next,

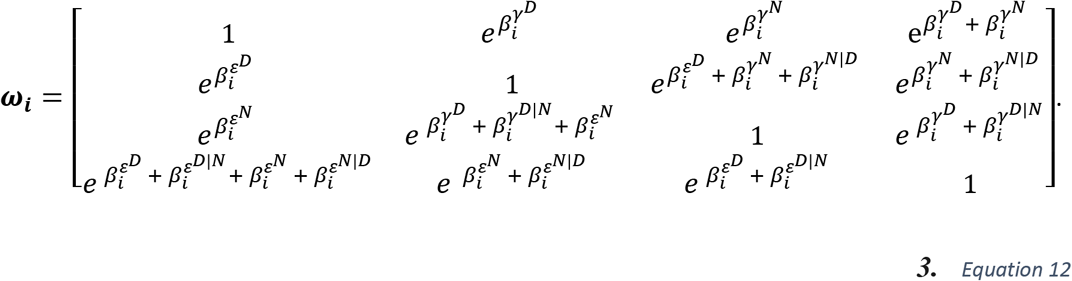

Dividing each element of a row by its respective row sum (i.e, applying the multinomial logit-link) converts ***ω_i_*** to ***τ_i_*** (Fidino et al. 2019). The reduced model removes all second-order parameters from ***ω_i_*** and becomes,

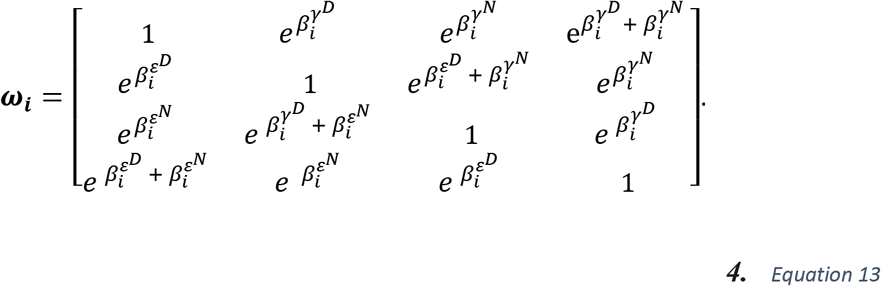

The null model, which is a multi-season occupancy model with covariates, ***ω_i_*** becomes,

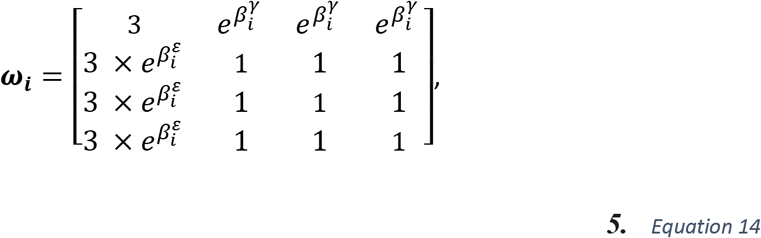

where 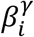 and 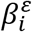 are respectively logit-linear predictors for colonization and extinction. The dynamic MSDOM with covariates uses the same process to incorporates detection-level covariates, save for the fact that the detection matrix and data vary across the secondary sampling periods.

###### Fosa Case Study

Fosa are a medium size carnivore (5.5-9.9 kg; Goodman 2012) in the monophyletic Eupleridae family, which is endemic to Madagascar. Fosa face increasing anthropogenic pressure from deforestation (Morelli et al., 2020), unsustainable hunting (Golden, 2009), and exotic species (Farris et al., 2017). As a generalist species with a diverse diet, fosa’s activity near human settlements and consumption of livestock has caused conflict with humans (Borgerson, 2016; Kotschwar Logan et al., 2014). Previous studies show their diel activity is largely cathemeral (Farris et al., 2015a; Gerber et al., 2012a). Their ubiquitous occurrence across forests and use of the entire 24-hour period make them an exemplar species to investigate the utility of MSDOM in the context of human disturbance. We analyzed data from Makira Natural Park (Farris et al., 2015b) and Ranomafana National Park regions (Gerber et al., 2012a; see Appendix S3; Table 1).

These two parks have unique histories which has shaped differing activity in each region (changes in forest cover, agriculture, invasive species introduction, etc.) and subsequent impact on native wildlife species (Goodman et al. 2019). As such, we have formed unique hypotheses about anthropogenic factors which impact these regions. Given high human activity within forests of Makira (Farris et al., 2015b) compared to Ranomafana (Farris et al., 2017; Gerber et al., 2012b), we used human activity at camera locations to quantify human disturbance. Human activity was calculated as the number of human detection events (photos taken within a 30 min intervals) per diel period (i.e., day and night) for each camera site divided by the number of sampling days the site was active. At Ranomafana, human activity within the protected boundaries were low in contrast to Makira. The riskiest areas for fosa at Ranomafana were found outside the park boundaries or along forest edges where villages are located and there is high human activity. Therefore, we used the distance to the nearest village and distance to the nearest matrix (non-forest) from each camera trap to quantify human disturbance (see Gerber et al. 2012a for details).

We fit static MSDOMs to the Makira and Ranomafana data separately. For both regions, we hypothesized that occupancy would vary in diel time by the level of disturbance. We also hypothesized the ‘day use’ state to be used least by fosa due to diurnal human activity near areas of high disturbance. Specifically, we predicted that fosa occupancy during the day would decrease with increasing human disturbance and fosa occupancy at night would be higher than day occupancy, regardless of human disturbance. We also expected increasing night occupancy with increasing human disturbance. Day was defined by hours after civil sunrise and before civil sunset, while night was defined by hours following civil sunset and before civil sunrise calculated through package suncalc (Thieurmel and Elmarhraoui, 2019) in R v 4.0.2. To determine detected diel states of fosa, we used 6-day occasions. All models were coded and fit in JAGS v. 4.0.2 (Plummer, 2003) with the runjags package (Denwood, 2016) in R v. 4.0.2. We assessed convergence using the Gelman–Rubin diagnostic (Gelman and Rubin, 1992) to ensure all values were < 1.1 and by visually examining traceplots of the posterior distributions. We compared models using the conditional predictive ordinate (CPO; Hooten and Hobbs, 2015) and evaluated evidence of an effect with the most supported model by investigating whether 95% credible intervals of parameter estimates included zero and deriving the probability of an effect being less than or greater than zero.

We fit 18 candidate models to the Ranomafana data: full, reduced, and null model, each with state-occupancy modeled with and without individual covariates (distance to village and matrix were modeled separately) and a categorical variable for survey (see Appendix S 3; Table 2). Detection parameters were not modeled with covariates. The most supported model was the full model with the covariate distance to village influencing state occupancy. We found strong support for 1) variation in state occurrence (Fig 1) and detection (see Appendix S3; Fig 1) and 2) multistate occurrence varying with human disturbance (Fig 2A). We found little support that day occurrence varied by distance to village based on the mode and 95% credible interval (α^Day,Dist.Vill^ = −0.002; 95% CI = −1.31, 1.37), with only a 0.50 probability that the distribution was above zero (Fig 1). This contrasts with our hypothesis. However, we found moderate to strong support that night-day occurrence increased with distance to village (α^ND,Dist.Vill^ = 1.45; 95% CI = −0.17, 2.95; P(α^ND,Dist.Vill^ > 0) = 0.97), supporting our hypothesis. These results suggest that if fosa use sites during day hours, it is in conjunction with night hours, and the probability of using sites during the day is greater further away from human disturbance. We also found moderate to strong support that night occurrence declined with increasing distance to village (α^Night,Dist.Vill^ = −1.16; 95% CI = −2.38, −0.02; P(α^Night,Dist.Vill^ < 0) = 0.98). Results from conditional probabilities of use (given fosa are present) revealed similar probabilities (Fig 2B) to those of occurrence. This was due to the widespread distribution of fosa within the study area. We found the probability to detect fosa at night, given it was present during the day and night (*p*^4,3^), to be the highest detection probability (see Appendix S3; Table 1). Detection of fosa during the day-night state (*p*^4,4^) was the lowest. This suggests that this low density and wide-ranging species does use sites during the day and night, but not regularly.

**Figure 1.**
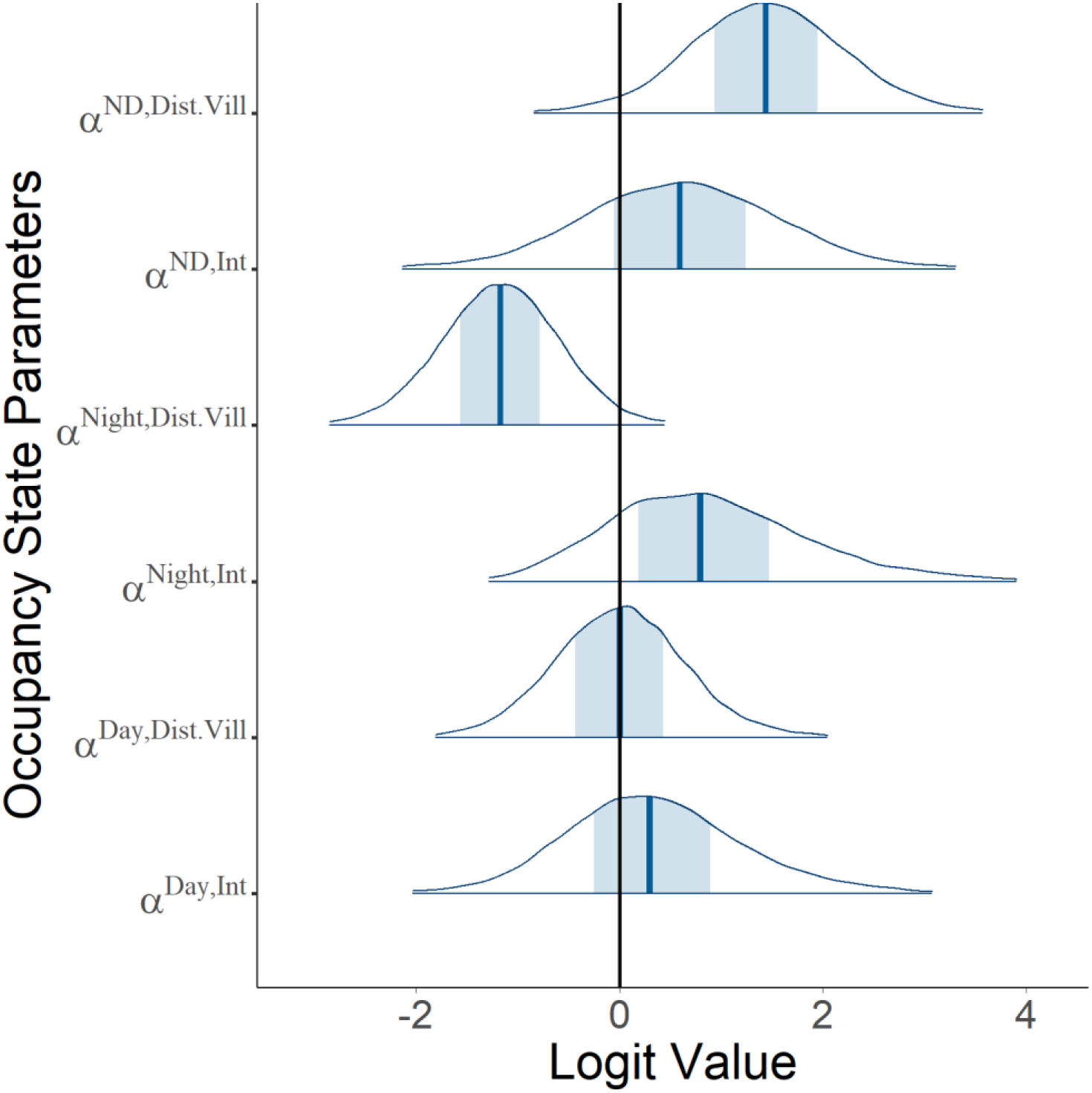
Posterior distributions of fosa (*Cryptoprocta ferox*) state occupancy model parameters for the most supported model using the Ranomafana National Park data. The light blue shaded area represents 50% probability density and the dark blue line indicates the posterior mode. Y-axis labels with “Int” indicate an intercept for the states (Day, Night, ND = Night and Day) and “Dist.Vill” indicates the slope parameter associated to the variable distance to village.

**Figure 2.**
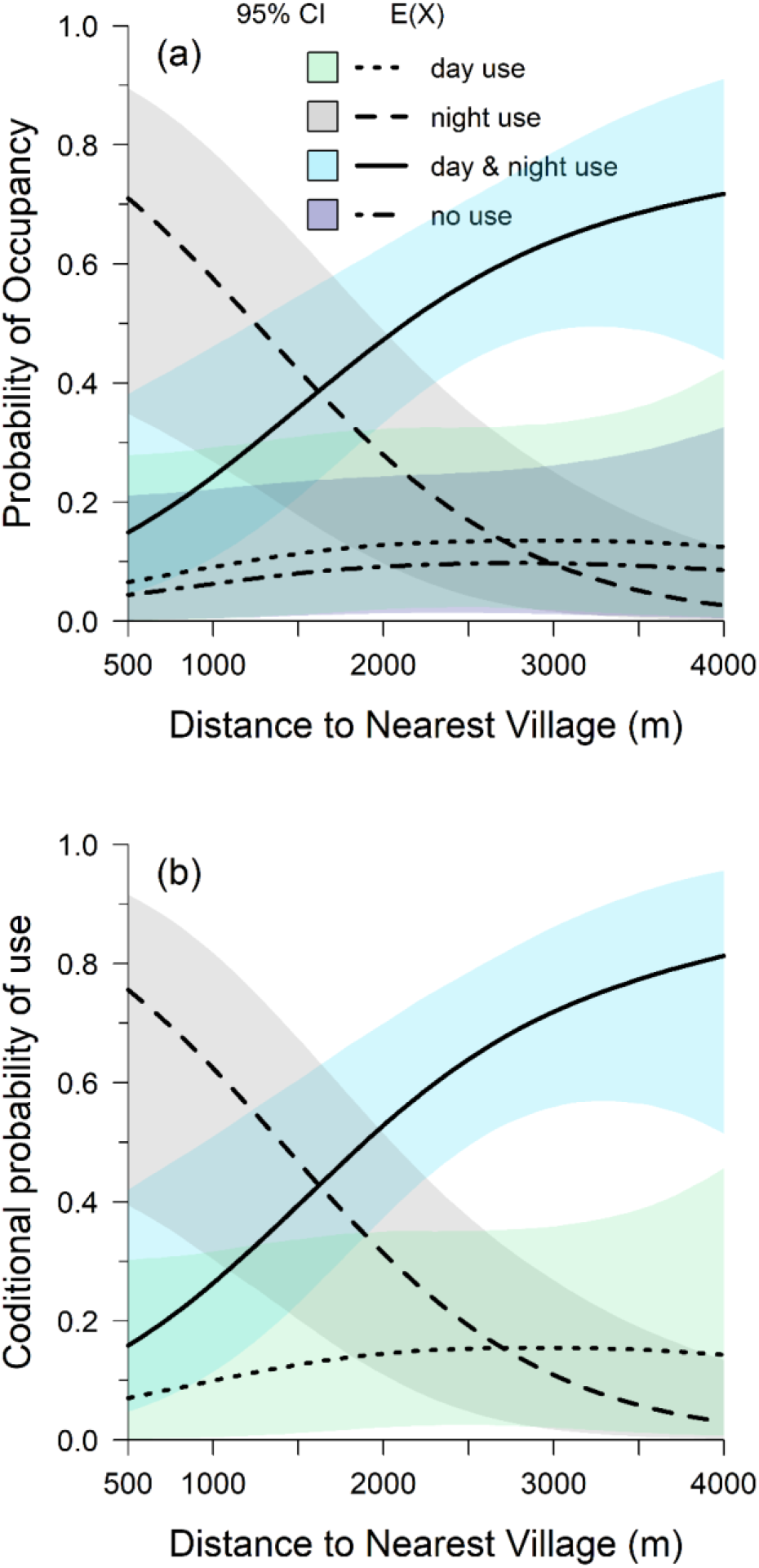
A) The probability of use for fosa (*Cryptoprocta ferox*) for each diel category with the expected value, E(X), the median of the posterior of a given parameter and 95% credible interval (CI), B) The probability of use for fosa at for each diel category given fosa are present with E(X) and 95% credible interval (note state 1, ‘no use’ is not included here). Both are estimated from the most supported model using the Ranomafana National Park data which incorporated the covariate distance to village.

We fit 6 candidate models with the Makira data: full, reduced, and null model, each with and without the human activity covariate; detection parameters were not modeled with covariates (see Appendix S3; Table 4). We found all models to fit the data (0.1>Bayesian GOF p-value<0.9). We found the most supported model to be the full model without an effect of human activity. These results support that there is variation in multistate occurrence and detection, but not regarding our hypothesis that human disturbance influenced occurrence. We found fosa occupancy was highest during the night state (*ψ*^3^ = 0.33; 95% CI = 0.11, 0.60), followed by day, (*ψ*^2^ = 0.20; 95% CI = 0.06, 0.44), and day & night state (**ψ**^4^ = 0.18; 95% CI = 0.05, 0.41; Fig 3). The large parametric uncertainty of the detection parameters made drawing conclusions difficult, though results indicate fosa are most detectable at night when present during the day & night state (see Appendix S3; Table 5).

**Figure 3.**
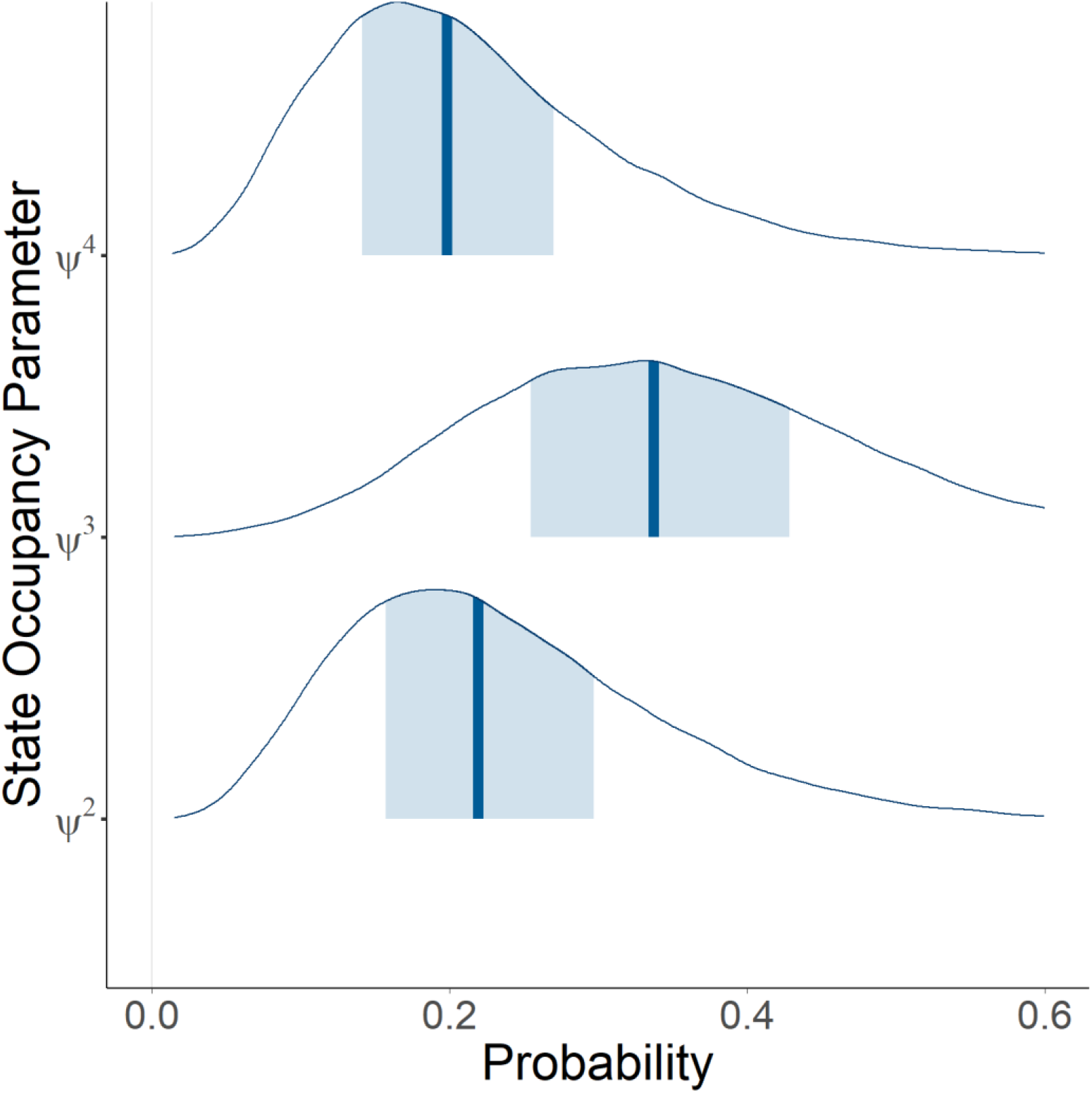
Posterior distributions of fosa (*Cryptoprocta ferox*) state occupancy model parameters (mean across all sites) for the most supported model using the Makira Natural Park data. The light blue shaded area represents 50% probability density and the dark blue line indicates the posterior mode. *ψ*^2^ is the probability a species is present during the day only; *ψ*^3^ is the probability a species is present during the night only; *ψ*^4^ is the probability a species is present during the day & night.

###### Coyote Case Study

Coyote are a medium sized carnivore (8-14 kg; Bekoff and Gese, 2003) native to North America that have expanded their distribution across the United States, Canada, and South America in the last century (Hody and Kays, 2018). As generalists, coyote exploit an array of habitats from prairies to urban cities (Elliot et al., 2016). Coyote diel activity is quite plastic, specifically in the presence of anthropogenic disturbance (Gehrt et al., 2007; Way et al., 2004). Therefore, we quantified whether coyote modify their diel activity along an urbanized gradient.

To do so, we fit dynamic MSDOMs to 13 sampling periods of camera trapping data collected between July 2016 and July 2019 in the greater Chicago Metropolitan area. Camera deployments followed protocols outlined by the Urban Wildlife Information Network (see Magle et al. 2019). Briefly, 105 cameras were placed along three 50 km transects radiating outward from downtown Chicago, Illinois, USA (see Appendix S3; Table 6). Data were summarized such that each 4-week deployment (e.g., July 2016, October 2016, etc.) was treated as a primary sampling period and each week was a secondary sampling period. To determine the detected diel state for a given week (occasion length), we used the suncalc package in R following the same diel categorization process as the fosa study. While the static MSDOM (with 4 states) can potentially have 3 linear predictors for the latent state, the dynamic MSDOM potentially has 11, thereby exacerbating the number of different covariate combinations and parameters to be estimated. To simplify our model fitting strategy, we fit 3 models that differed in their fundamental structure (i.e., the full, reduced, and null dynamic MSDOM), and included an urban intensity metric on all first-order parameters. We made two additional changes to the full model because daytime coyote detections were sparse (n = 54) relative to night (n = 286) or day & night (n = 183). First, we excluded urban intensity on second-order colonization or extinction parameters because second-order slope terms failed to converge when included. Second, we used Eq. 3 as the detection matrix, which assumes that the probability of detecting ‘day & night use’ (state 4) as the product of the probabilities of detecting ‘day use’ (state 2) and ‘night use’ (state 3). Models were compared with CPO and we evaluated evidence of an effect with the best-fit model by investigating whether 95% credible intervals of parameter estimates included zero and deriving the probability of an effect being less than or greater than zero.

To derive the urban intensity metric, we used principal component analysis for tree cover (%; CMAP, 2016), impervious cover (%; CMAP, 2016), and housing density (units km^-2^; Hammer et al. 2004) within a 1-km buffer of each sampling location. Negative values represented increased forest cover coupled with decreased impervious cover and housing density, while positive values represented increased levels of impervious cover and housing density coupled with low canopy cover. Models were fit in JAGS v 4.3.0 in R v 4.0.3. We evaluated model convergence by inspecting traceplots to ensure proper mixing and using the Gelman-Rubin statistic.

Of the possible 1365 deployments (105 sites across 13 sampling periods), we collected data for 1172 deployments. ‘No use’ was the most observed state (n = 650), followed by ‘night use’ (n = 286), ‘day & night use’ (n = 183), and ‘day use’ (n = 53). Overall, the full model (22 parameters, CPO = 3131.46) had the most support, followed by the reduced (16 parameters, CPO = 3209.17) and then the null model (8 parameters, CPO = 3334.52). With the most supported model, the average occupancy probability during the first season was 0.41 for ‘no use’ (95% CI = 0.26, 0.56), 0.18 for ‘day use’ (95% CI = 0.06, 0.33), 0.07 for ‘night use’ (95% CI = 0.01, 0.19), and 0.32 for ‘day & night use’ (95% CI = 0.19, 0.48). Thus, assuming a site was occupied by coyote during the first primary period, coyote were on average, most likely to use sites during the day and night. Across the urbanization gradient, ‘day use’ was more negatively associated to urban intensity (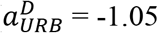, 95% CI = −1.98, −0.07, 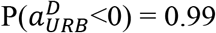) than ‘night use’ (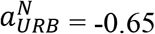, 95% CI = −1.51, 0.18, 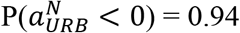). There was some evidence that ‘day & night’ use became more common with increasing urban intensity, but 95% credible intervals for this second-order parameter overlapped 0 (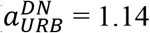, 95% CI = −0.08, 2.50, 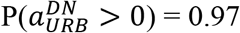). While the initial occupancy parameters demonstrate that ‘day use’ decreases with increasing levels of urban intensity, it is only a snapshot of the underlying process. The dynamic MSDOM provides new ways to assess this relationship through additional manipulations of the latent-state transition probability matrix (**τ)**, which describe the processes that bring about coyote occupancy.

While it is equally important to explore the underlying colonization and extinction dynamics of the model, by solving the equation ***δ**_i_* = ***δ**_i_**τ**_i_* where **∑ *δ***_*i*_ = 1 it is possible to derive the expected probability of each occupancy state at each site (Fidino et al. 2019). Doing so simplifies the *I* × M × M transition matrix into *I* × M occupancy probabilities, and therefore can highlight the overall pattern across an environmental gradient. We applied this equation to the entire posterior of ***τ**_i,t_*, and generated predicted occupancy states at hypothetical sites across Chicago’s urbanization gradient. Following this, the probability of use of the different coyote occupancy states, conditional on coyote presence, can be derived by calculating the conditional probability of ‘day use’, ‘night use’ and ‘day & night’ given coyote presence. For example, 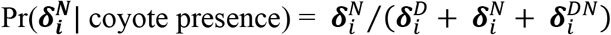. Plotting these relationships reveals that while ‘day & night use’ is the most likely category at low levels of urban intensity, it is replaced by ‘night’ use as urban intensity increases, assuming coyote are present (Fig. 4).

**Figure 4.**
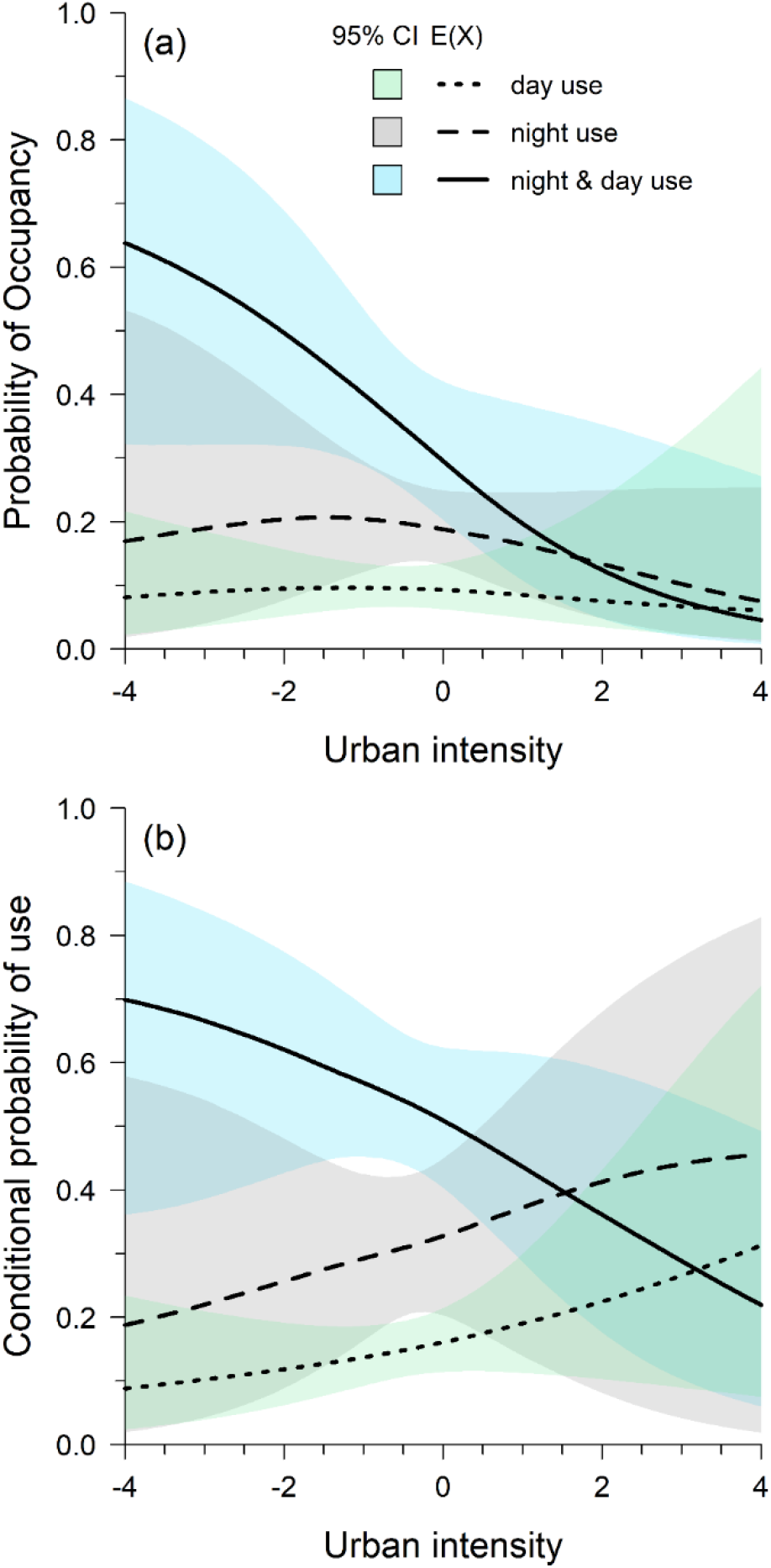
A) The probability of use for coyote (*Canis latrans*) each diel category with the expected value, E(X), the median of the posterior of a given parameter and 95% credible interval (CI; note state 1, ‘no use’ not included here), B) The probability of use for each diel category given coyote presence with E(X) and 95% credible interval (note state 1, ‘no use’ not included here). Both are estimated from 13 sampling periods of camera trapping data from 1172 deployments at 105 unique sites between July 2016 and July 2019. In both cases. as urban intensity increases the ‘night use’ diel category becomes the most common category, though there is uncertainty in this estimate given the credible intervals.

The transitions among different states can be plotted out and interpreted through the parameters that describe them (Fig. 5). For example, sites without coyotes were most likely to stay in the ‘no use’ state across all levels of urban intensity, though this relationship became more pronounced at high levels of urban intensity (Fig. 5). The transitions from ‘no use’, which are described by γ^*D*^ and γ^*N*^, were driven by the strongly negative first-order colonization intercepts for ‘day use’ (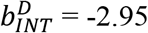, 95% CI = −3.88, −2.14, 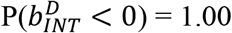) and night use (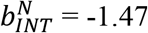, 95% CI = −1.85, −1.10, 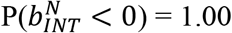), as well as a negative association between ‘night use’ and urban intensity (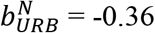, 95% CI = −0.62, −0.09, 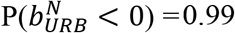). There was weak support that colonization of ‘day use’ negatively covaried with urban intensity (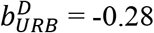, 95% CI = −0.74, 0.16, 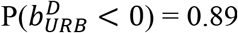). While ‘night use’ negatively covaried with urban intensity, the relatively less negative intercept of this level of the model (i.e., 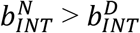) made ‘night use’ the most likely diel category for coyotes to colonize along the gradient of urban intensity (Fig. 5).

**Figure 5.**
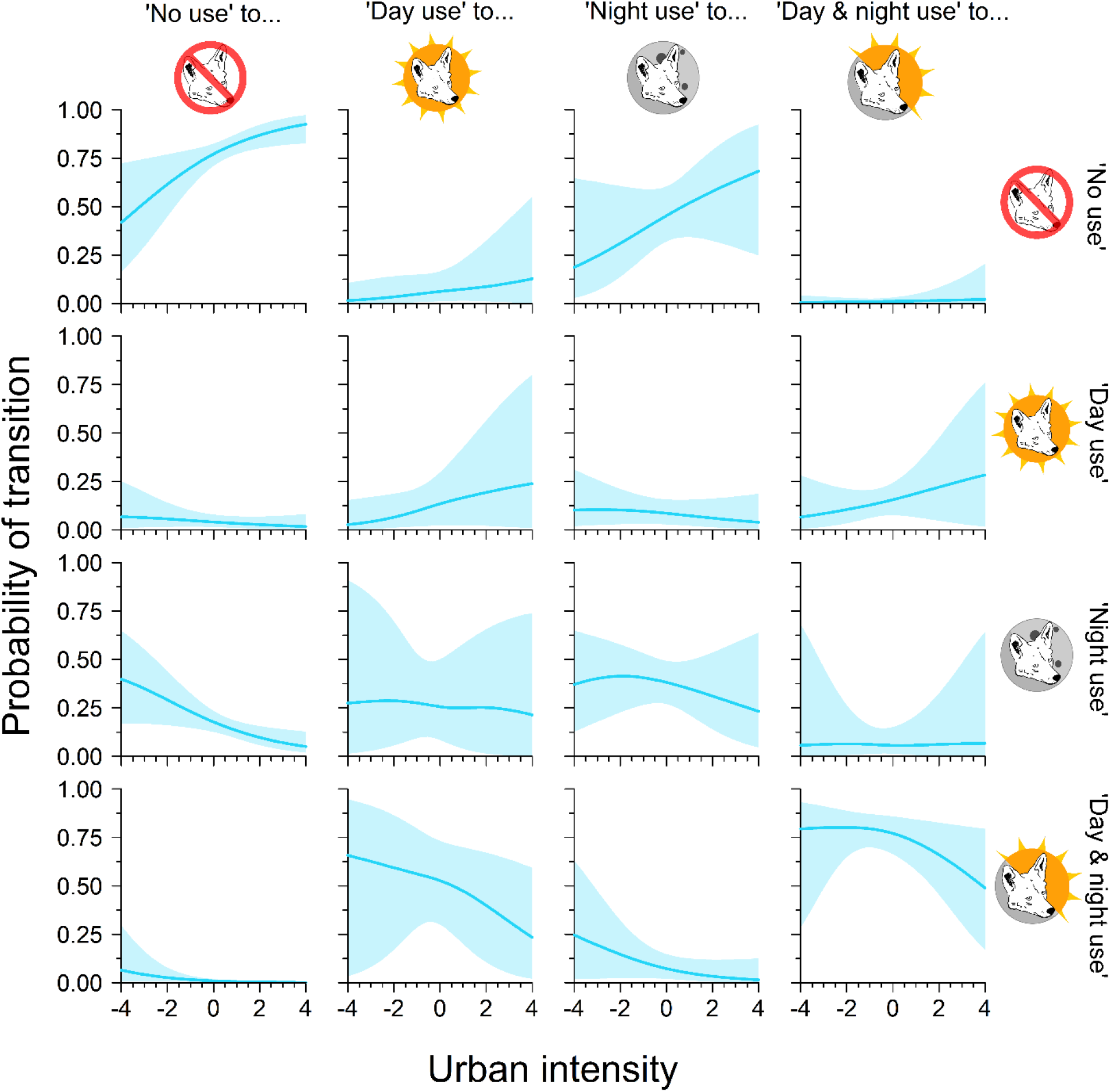
The transition probabilities among each of the four diel coyote (*Canis latrans*) states as a function of urban intensity estimated from 13 sampling periods of camera trapping data collected in Chicago, Illinois, USA from 1172 deployments at 105 unique sites between July 2016 and July 2019. Horizontal lines indicate the median estimate while shaded ribbons are 95% credible intervals.

When a site was in the ‘night use’ state, transitions are described by *ε^N^* and *γ*^*D|N*^. At average levels of urban intensity, sites were most likely to transition to ‘day & night use’ (0.53, 95% CI = 0.30, 0.73), followed by ‘night use’ (0.26, 95% CI = 0.08, 0.50), ‘day use’ (0.13, 95% CI = 0.02, 0.30), and then ‘no use’ (0.06, 95% CI = 0.01, 0.17). The large increase in ‘day & night use’ was driven by the positive second-order ‘night use’ colonization parameter (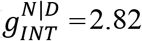, 95% CI = 1.60, 4.52, 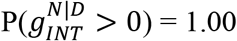), whereas the decreasing transition probability of ‘day use’ to ‘day & night use’ was governed by the negative first-order ‘night use’ colonization slope term (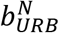, listed above). Likewise, first-order ‘day use’ extinction rates were relatively modest (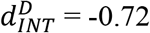, 95% CI = −2.02, 0.43, 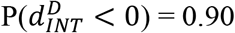) and covaried little with urban intensity (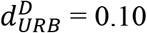, 95% CI −1.04, 0.85, 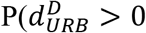). As a result, 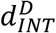 and 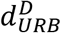 generated relatively flat transitions from ‘day use’ to either ‘no use’, ‘night use’, or back to ‘day use’ (Fig. 5).

Finally, at ‘day & night use’, transitions are described by ε^*D*|*N*^ and ε^*N*|*D*^. Second-order parameters associated to these probabilities were both strongly negative (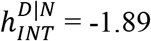, 95% CI = −3.67, −0.17, 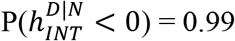, 95% CI = −2.70, −0.98, 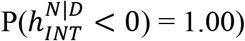). When these second-order parameters are combined with the relatively small influence urban intensity had on first-order extinction parameters (i.e., 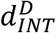 and 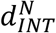), sites in ‘day & night’ use were by far more likely to remain in this state (Fig. 5).

In regard to detectability, if a site was in state ‘day use’ the probability of detecting that state was 0.15 (95% CI = 0.12, 0.18) at average levels of urban intensity, and covaried little with urban intensity (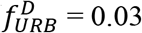, 95% CI = −0.20, 0.22, 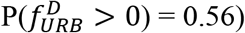). The ability to detect ‘night use’ was, on average, double that of ‘day use’ (0.30, 95% CI = 0.28, 0.33), but was minimally and negatively associated to urban intensity (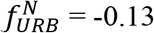, 95% CI = −0.24, −0.01, 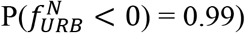). When a site was in ‘day & night’ use, at average levels of urban intensity we were most likely to observe the site as ‘no use’ (0.59, 95% CI = 0.56, 0.62), followed by ‘night use’ (0.26, 95% CI = 0.23, 0.28), ‘day use’ (0.10, 95% CI = 0.09, 0.12), then ‘day & night use’ (0.04, 95% CI = 0.04, 0.05).

## 6. DISCUSSION

The study of animal-habitat relationships has often focused on identifying spatial drivers of species occurrence, while largely ignoring *when* species use habitat within the diel period. We developed the MSDOM framework to allow species’ spatial habitat use to be studied over a fine (diel) temporal scale within and across seasons or years. Importantly, our framework allows detection probability to vary across sites used at different diel periods, a source of heterogeneity that is typically unmodeled and is required to produce unbiased parameter estimates. The utility of the MSDOM framework is especially pertinent to studying species at risk to human activities and easily accommodates continuous measures of anthropogenic change on diel behavior. This is important because one way species adapt to changing landcover or human development is to shift their activity away from diel periods of high risk (Gaynor et al., 2018; Gaston, 2019). Such behavioral shifts are likely not without important ecological costs and may go undetected under previous models, but can be detected with the MSDOM.

Our case studies highlight that spatial habitat is not used equally across diel time. We found that fosa and coyote temporally structure their site-use in response to anthropogenic drivers. Previous studies of fosa in the eastern rainforests have suggested that they are ubiquitously distributed across forested landscapes and are predominantly cathemeral (Gerber et al. 2012b; Farris et al 2015a). By jointly investigating fosa spatial-temporal habitat use, we found that they do occur widely across forested sites, but vary when they use a site based on its proximity to anthropogenic activity. For example, fosa at Ranomafana were nocturnal near human villages, which occur along the edges of the protected forest. At the forest interior, fosa were cathemeral. These findings indicate that within specific habitats, fosa can be active during day and night hours, but human activity and development limit fosa to roughly half of their potential activity period. However, the level and type of human disturbance is important in predicting fosa diel activity, as we did not find support that human activity affected diel occurrence at Makira; this is likely due to predictable diurnal human activity and locations of camera sites which were connected to core forest habitat at greater distances to human villages (Farris et al 2015b).

Like fosa, we found variation in coyote diel activity across anthropogenic gradients. Overall, we found that coyote used sites during the day and night at low levels of urban intensity, which likely reflects that this species is crepuscular in natural environments (McClennen et al. 2001). However, as urban intensity increased, diel use of sites transitioned to be nocturnal. In combination with this, we found that the marginal occupancy of coyote, irrespective of diel state, decreased with increasing urban intensity. Thus, while coyote occupy less habitat in the core of Chicago, the habitat they do occupy is generally used at night.

A special feature of the dynamic MSDOM is that the transition matrix provides additional information on diel use which helps disentangle the expected occupancy patterns in how coyote used diel time across space. For example, while it was relatively rare for coyotes to use highly urban sites during the day and night, their probability to persist from one season to the next in this state was high. Conversely, coyote were most likely to use highly urban sites only at night, but were most likely to go locally extinct when this occurred (i.e., transition to state “no use”). Thus, even though coyote were more likely to use highly urban sites at night, the use of these sites is more ephemeral than the urban sites coyote use throughout the entire diel period. Because urban coyotes typically have home ranges roughly twice the size of their rural counterparts (Gese et al., 2012), we suspect that in the urban core coyote use pockets of primary habitat during the day and night and venture out to secondary or tertiary habitat patches exclusively at night, when human activity levels are low.

As the definition of habitat evolved to better recognize the value of time, so too should our modeling approaches. Our MSDOM achieves this and can measure the effect of continuous covariates to quantify change in diel behavior across space and though time. Although understanding habitat use of species has been critical in making informed conservation and management decisions (Guisan et al., 2013), current land-planning tools are often limited to spatial considerations (Gaynor et al., 2018). Though progress has been made in protecting habitats used over longer timescales, such as seasons, we lacked informative tools to protect habitat during critical diel periods. Advanced modeling approaches that estimate diel-habitat use will be a valuable asset in supporting successful conservation and land-management strategies in a rapidly changing world.

## ACKNOWLEDGEMENTS

This work was supported by the USDA National Institute of Food and Agriculture, Hatch Formula project 1017848, the Prentice Foundation, and the Davee Foundation. We are grateful to S. Karpanty and M. Kelly for their contributions to the fosa studies and to Institute of Conservation of Tropical Ecosystems (MICET) and permission and support from the Government of Madagascar and Madagascar National Park Services.

