## Appendix S1 for "Rethinking habitat occupancy modeling and the role of diel activity in an anthropogenic world" (from another version of this preprint: version 1, 2021-07-01)

### Appendix S1: Additional data collection details and methodology

| Parameter type | Verbal description |
|---|---|
| $\boldsymbol{\psi}$ | Probability of occupancy vector for the *M* = 4 states, which sums to one. The full model assumes that state 4 ('day & night') is independent whereas the reduced model assumes state 4 is the product of states 2 ('day') and 3('night', Eq. 2). The null model assumes states 2, 3, and 4 are all equal (Eq. 4). For all models, states 2, 3, and 4 are estimated from the data while state 1 is obtained through subtraction such that $\psi^1 = 1 - \psi^2 - \psi^3 - \psi^4$. |
| $\boldsymbol{P}$ | Probability of detection matrix for the M = 4 states, which is $M \times M$ with the observed states along the column and true states along the rows. Rows sum to one. The full model assumes that there is no relationship among state-specific detection probabilities, and so detection probabilities in state 4 ('day & night') are not defined by the detection probabilities of state 2('day') or 3 ('night', Eq. 1). Conversely, the reduced model assumes that the detection probabilities in state 4 are related to states 2 and 3, or their complementary probability, depending on the observed state (Eq. 3). The null model assumes the probability of detecting states 2, 3, or 4 are identical (eq. 5). |
| $\boldsymbol{\tau}$ | The $M \times$ M transition probability matrix of the dynamic MSDOM. The true state at sampling period *t*-1 is along the rows and the possible states to transition to are along the columns. Each row sums to 1. Transitions are composed of local colonization ($\gamma$) and extinction (ε) probabilities. To reduce model complexity, the full (Eq. 8), reduced (Eq. 9), and null (Eq. 10) models assume all transitions are defined by the product of local colonization and extinction probabilities, though they do so in different ways (described below). |
| $\boldsymbol{\gamma}$ | The probability of local colonization between sampling period *t*-1 and *t*. The full model (Eq. 8) has four colonization probabilities: the probability an unoccupied site transitions to state 2 ('day', $\gamma^D$) or state 3('night', $\gamma^N$), the conditional probability a site transitions to state 2 given state 3 at *t*-1 ($\gamma^{D\|N}$), and the conditional probability a site transitions to state 3 given state 2 at *t*-1 ($\gamma^{N\|D}$). The reduced model (Eq. 9), removes the conditional probabilities present in the full model and therefore assumes that transitions between day and night are composed of $\gamma^D$ and $\gamma^N$. The null model (Eq. 10) assumes day and night transitions are the same so there is a single colonization probability ($\gamma$). |
| $\boldsymbol{\varepsilon}$ | The probability a local extinction between sampling period *t*-1 and *t*. The full model (Eq. 8), has four extinction probabilities: the probability state 2 ('day') transitions to any other state ($\varepsilon^D$), the probability state 3 ('night') transitions to any other state ($\varepsilon^N$), the conditional probability day is not included at time *t* given state 4 ('day & night') at time *t*-1 ($\varepsilon^{D\|N}$), and the conditional probability night is not included at time *t* given state 4 at time *t*-1 ($\varepsilon^{N\|D}$). The reduced model (Eq. 9) removed the conditional probabilities present in the full model and therefore assumes that transitions between day and night are composed of $\varepsilon^D$ and $\varepsilon^N$. The null model (Eq. 10) assumes day and night transitions are the same so there is a single extinction probability (ε). |

Table 1. A high-level description of the types of model parameters for the multi-state diel occupancy model (MSDOM). For this description, we assume that there are *M* = 4 states which represent whether the species is 1) not present, 2) present during the day only, 3) present during the night only, and 4) present during the day & night.

| **Parameters** | **Relevant Model** | **Verbal Description** |
|---|---|---|
| $p^{2,2} = p^{\mathrm{DAY}}$ | Full | Probability of detecting a species during the day only, given it was present only during the day. Part of matrix P. |
| $p^{3,3} = p^{\mathrm{Night}}$ | Full | Probability of detecting a species at night only, given it was present only at night. Part of matrix P. |
| $p^{4,1} = p^{\mathrm{ND1}}$ | Full | Probability of not detecting a species, given it was present during the day and night. Part of matrix P. |
| $p^{4,2} = p^{\mathrm{ND2}}$ | Full | Probability of detecting a species during the day only, given it was present during the day and night. Part of matrix P. |
| $p^{4,3} = p^{\mathrm{ND3}}$ | Full | Probability of detecting a species during the night only, given it was present during the day and night. Part of matrix P. |
| $p^{4,4} = p^{\mathrm{ND4}}$ | Full | Probability of detecting a species during the day and night, given it was present during the day and night. Part of matrix P. |
| $p^{\mathrm{Day.M}}$ | Reduced | Probability of detecting a species during the day, regardless of use or lacking use at night. Part of matrix P. |
| $p^{\mathrm{Night.M}}$ | Reduced | Probability of detection a species at night, regardless of use or lacking use during the day. Part of matrix P. |
| $\psi^{\mathrm{Day.M}}$ | Reduced | Probability of using a site during the day, regardless of use or lacking use at night. Part of probability of occupancy vector. |
| $\psi^{\mathrm{Night.M}}$ | Reduced | Probability of using a site at night, regardless of use or lacking use during the day. Part of probability of occupancy vector. |
| $\psi^1$ | Full, Reduced, Null | Probability a species is not present. Part of probability of occupancy vector. |
| $\psi^2$ | Full, Reduced, Null | Probability a species is present during the day only. Part of probability of occupancy vector. |
| $\psi^3$ | Full, Reduced, Null | Probability a species is present during the night only. Part of probability of occupancy vector. |

| | | |
|---|---|---|
| $\psi^4$ | Full, Reduced, Null | Probability a species is present during the day & night. Part of probability of occupancy vector. |
| $\psi^{\bullet}$ | Full, Reduced, Null | The probability a site is occupied, regardless of state. Part of probability of occupancy vector. |
| $p^{\bullet}$ | Null | The probability of detection, regardless of state. Part of matrix P. |

Table 2. Verbal descriptions and model relevancy of fundamental model parameters for the static (single season) multi-state diel occupancy model.

| **Parameters** | **Relevant Model** | **Verbal Description** |
|---|---|---|
| $p_t^{2,2} = p_t^{\text{DAY}}$ | Full | Probability of detecting a species in season *t* during the day only, given it was present only during the day. Part of matrix P. |
| $p_t^{3,3} = p_t^{\text{Night}}$ | Full | Probability of detecting a species in season *t* at night only, given it was present only at night. Part of matrix P. |
| $p_t^{4,1} = p_t^{\text{ND1}}$ | Full | Probability of not detecting a species in season *t*, given it was present during the day and night. Part of matrix P. |
| $p_t^{4,2} = p_t^{\text{ND2}}$ | Full | Probability of detecting a species in season *t*, during the day only, given it was present during the day and night. Part of matrix P. |
| $p_t^{4,3} = p_t^{\text{ND3}}$ | Full | Probability of detecting a species in season *t*, during the night only, given it was present during the day and night. Part of matrix P. |
| $p_t^{4,4} = p_t^{\text{ND4}}$ | Full | Probability of detecting a species in season *t*, during the day and night, given it was present during the day and night. Part of matrix P. |
| $p_t^{\text{Day.M}}$ | Reduced | Probability of detecting a species in season *t*, during the day, regardless of use or lacking use at night. Part of matrix P. |
| $p_t^{\text{Night.M}}$ | Reduced | Probability of detection a species in season *t*, at night, regardless of use or lacking use during the day. Part of matrix P. |
| $\psi_1^{\text{Day.M}}$ | Reduced | Probability of using a site in season 1, during the day, regardless of use or lacking use at night. Part of probability of occupancy vector. |
| $\psi_1^{\text{Night.M}}$ | Reduced | Probability of using a site in season 1, at night, regardless of use or lacking use during the day. Part of probability of occupancy vector. |
| $\psi_1^1$ | Full, Reduced, Null | Probability a species is not present in season 1. Part of probability of occupancy vector. |
| $\psi_1^2$ | Full, Reduced, Null | Probability a species is present during the day only in season 1. Part of probability of occupancy vector. |

| | | |
|---|---|---|
| $\psi_1^3$ | Full, Reduced, Null | Probability a species is present during the night only in season 1. Part of probability of occupancy vector. |
| $\psi_1^4$ | Full, Reduced, Null | Probability a species is present during the day & night in season 1. Part of probability of occupancy vector. |
| $\psi^{\bullet}$ | Full, Reduced, Null | The probability a site is occupied, regardless of state. Part of probability of occupancy vector. |
| $p^{\bullet}$ | Null | The probability of detection, regardless of state. Part of matrix P. |
| $\gamma^D$ | Full, Reduced | The probability a site is colonized and used in the day only. Part of the probability of local colonization between sampling period *t*-1 and *t*. |
| $\gamma^N$ | Full, Reduced | The probability a site is colonized and used in the night only. Part of the probability of local colonization between sampling period *t*-1 and *t*. |
| $\gamma^{N\|D}$ | Full | The probability a site is colonized and used in the night only, given it was used previously in the day only. Part of the probability of local colonization between sampling period *t*-1 and *t*. |
| $\gamma^{D\|N}$ | Full | The probability a site is colonized and used in the day only, given it was used previously in the night only. Part of the probability of local colonization between sampling period *t*-1 and *t*. |
| $\varepsilon^D$ | Full, Reduced | The probability a site is no longer used in the day only. Part of the probability of local extinction between sampling period *t*-1 and *t*. |
| $\varepsilon^N$ | Full, Reduced | The probability a site is no longer used in the night only. Part of the probability of local extinction between sampling period *t*-1 and *t*. |
| $\varepsilon^{N\|D}$ | Full | The probability a site is no longer used in the night only, given it was used previously in the day only. Part of the probability of local extinction between sampling period *t*-1 and *t*. |

| | | |
|---|---|---|
| $\varepsilon^{D\|N}$ | Full | The probability a site is no longer used in the day only, given it was used previously in the night only. Part of the probability of local extinction between sampling period *t*-1 and *t*. |

Table 3. Verbal descriptions and model relevancy of fundamental model parameters for the dynamic (multi-season) multi-state diel occupancy model. Descriptions also include matrix, vector, and probability affiliations described in Table 1.

| **Derived Parameter** | **Relevant Model** | **Verbal Description** |
|---|---|---|
| $\psi^4/(\psi^2 \times \psi^3)$ | Full Static and Dynamic | This day-night interaction factor quantifies the tendency for a site to be used during the day and night more than (>1) or less (<1) than expected; this will be exactly 1 in the reduced model, as it's part of the assumption of the model. This parameter is akin to a species interaction factor of co-occurrence models (MacKenzie et al., 2004). |
| $\frac{\psi^2}{\psi^\bullet}$ | Full and Reduced Static and Dynamic | The probability of temporal use at a site during the day only, given the site is used. |
| $\frac{\psi^3}{\psi^\bullet}$ | Full and Reduced Static and Dynamic | The probability of temporal use during the night only, given the site is used. |
| $\frac{\psi^4}{\psi^\bullet}$ | Full and Reduced Static and Dynamic | The probability of temporal use during the day & night, given the site is used. |

Table 4. Verbal descriptions and model relevancy of derived model parameters for the multi-state diel occupancy model.
