## Appendix S2 for "Rethinking habitat occupancy modeling and the role of diel activity in an anthropogenic world" (from another version of this preprint: version 1, 2021-07-01)

#### Appendix S2: Static modeling equations and methodology

### 1 Single Season MSDOM

#### 1.1 Fully State Specific

##### 1.1.1 Probability Model

We assume each $i = 1...N$ sites (spatial locations) are sampled with $j = 1....K$ independent surveys. Each site is considered to be in one of $M$ mutually exclusive states during the survey period in which we can assume there are no changes in the state occurrence (i.e., closure). We will define four total states throughout ($M = 4$) where the states of site $i$ are 1) not occupied, 2) occupied during the day only, 3) occupied during the night only, and 4) occupied during the night and day. State occurrence probabilities in the same order are,

$$\boldsymbol{\psi} = \begin{bmatrix} \psi^1 & \psi^2 & \psi^3 & \psi^4 \end{bmatrix}, \tag{1}$$

where $\psi^1 = 1 - \psi^2 - \psi^3 - \psi^4$ and $\mathbf{1} \cdot \boldsymbol{\psi} = 1$. The overall marginal (regardless of the state) occurrence of the species is $\psi^{\bullet} = \psi^2 + \psi^3 + \psi^4$ The true occupancy state for site $i$ is defined by the latent variable,

$$\mathbf{z}_i \sim \text{Categorical}(\boldsymbol{\psi}) \tag{2}$$

and is a 1 x $M$ matrix. For each true state, there is an order to which states may be observed, but this is not strictly hierarchical, as commonly applied in multi-state occupancy models (Nichols et al., 2007). Specifically, if a site is in state 3 (night only), it can not be observed in state 2 (day only). Let, $p_j^{m,l}$ be the probability of observing the occupancy state $l$, given the true state is $m$ in survey $j$. To maintain the nomenclature of our intended inference and assuming no site- or survey-level variation in detection probabilities, we can define $p^{2,2}$ as the probability of detection during the day state ($p^{\text{Day}}$), $p^{3,3}$ as the probability of detection during the night state ($p^{\text{Night}}$), and the detection probability in state 4 (in order) as $pND^1$, $pND^2$, $pND^3$, and $pND^4$. The detection matrix for survey $j$ is $M$ x $M$ with the observed (columns) and true state (row) as,

$$\boldsymbol{P} = \begin{bmatrix} 1 & 0 & 0 & 0 \\ 1 - p^{\text{Day}} & p^{\text{Day}} & 0 & 0 \\ 1 - p^{\text{Night}} & 0 & p^{\text{Night}} & 0 \\ p^{\text{ND1}} & p^{\text{ND2}} & p^{\text{ND3}} & p^{\text{ND4}} \end{bmatrix} \tag{3}$$

where $p^{1,4} = p^{\text{ND1}} = 1 - pND^2 - pND^3 - pND^4$ and $\boldsymbol{P} \cdot \mathbf{1} = 1$. The observed state at site $i$ in survey $j$ is a 1 x $M$ categorical random variable as,

$$\mathbf{y}_{ij} \sim \text{Categorical}(\boldsymbol{P}_{\mathbf{z}_i,}). \tag{4}$$

Diffuse prior distributions could be defined as,

$$
\begin{aligned}
\boldsymbol{\psi} &\sim \text{Dirichlet}(1,1,1,1) \\
p^{2,2} = p^{\text{Day}} &\sim \text{Beta}(1,1) \\
p^{3,3} = p_j^{\text{Night}} &\sim \text{Beta}(1,1) \\
\boldsymbol{p}^{4\bullet} = \boldsymbol{p}^{\text{ND}\bullet} &\sim \text{Dirichlet}(1,1,1,1).
\end{aligned}
$$

Under this model, there is no shared relationship between the state occupancy probability of state 4 (night and day) and states 2 (day only) and 3 (night only). There is also no relationship shared between the detection probabilities of the same states. This model would suggest that the detection or availability of the species at state 4 is possibly different then the detection in other states. This might be because of the species' behavior at sites used throughout the diel period is different than at the other sites. This model is coded for JAGS in the file 'jags.multistate.occ.full.R'.

##### 1.1.2 Multinomial-logit Model 1

It is common for model optimization reasons, the desire to use certain prior distributions, or when using covarites to model probabilities to recast occupancy and detection probabilities using a multinomial logit link as,

$$
\begin{aligned}
\mathbf{z}_i &\sim \text{Categorical}(\boldsymbol{\psi}) \\
\boldsymbol{\psi} &= \begin{bmatrix} \psi_i^1 & \psi_i^2 & \psi_i^3 & \psi_i^4 \end{bmatrix}. \\
\psi^1 &= \frac{\phi_i^1}{\phi_i^1 + \phi_i^2 + \phi_i^3 + \phi_i^4} \\
\psi^2 &= \frac{\phi_i^2}{\phi_i^1 + \phi_i^2 + \phi_i^3 + \phi_i^4} \\
\psi^3 &= \frac{\phi_i^3}{\phi_i^1 + \phi_i^2 + \phi_i^3 + \phi_i^4} \\
\psi^4 &= \frac{\phi_i^4}{\phi_i^1 + \phi_i^2 + \phi_i^3 + \phi_i^4}. \\
\phi^1 &= 1 \\
\phi^2 &= e^{\alpha_1} \\
\phi^3 &= e^{\alpha_2} \\
\phi^4 &= e^{\alpha_1+\alpha_2+\alpha_3}
\end{aligned}
$$

Further, we could do the same for the detection parameters as,

$$
\boldsymbol{q} = \begin{bmatrix} 1 & 0 & 0 & 0 \\ 1 & q^{2,2} & 0 & 0 \\ 1 & 0 & q^{3,3} & 0 \\ 1 & q^{4,2} & q^{4,3} & q^{4,4} \end{bmatrix}
$$

$$
\begin{aligned}
q^{2,2} &= e^{\beta^{\text{Day}}} \\
q^{3,3} &= e^{\beta^{\text{Night}}} \\
q^{4,2} &= e^{\beta^{\text{ND2}}} \\
q^{4,3} &= e^{\beta^{\text{ND3}}} \\
q^{4,4} &= e^{\beta^{\text{ND2}}+\beta^{\text{ND3}}+\beta^{\text{ND4}}}
\end{aligned}
$$

and the probability of detection parameters can be derived as,

$$
\begin{aligned}
p^{\text{Day}} &= \frac{q^{2,2}}{1 + q^{2,2} + 0 + 0} \\
p^{\text{Night}} &= \frac{q^{3,3}}{1 + 0 + q^{3,3} + 0} \\
p^{\text{ND1}} &= \frac{1}{1 + q^{4,2} + q^{4,3} + q^{4,4}} \\
p^{\text{ND2}} &= \frac{q^{4,2}}{1 + q^{4,2} + q^{4,3} + q^{4,4}} \\
p^{\text{ND3}} &= \frac{q^{4,3}}{1 + q^{4,2} + q^{4,3} + q^{4,4}} \\
p^{\text{ND4}} &= \frac{q^{4,4}}{1 + q^{4,2} + q^{4,3} + q^{4,4}}.
\end{aligned}
$$

A diffuse prior for all coefficients could be done using Logistic$(0, 1)$. This model is coded for JAGS in the file 'jags.multistate.occ.full.alt.R'.

##### 1.1.3 Multinomial-logit Model 2

To model site-level occurrence according to a single continuous covariates ($\mathbf{x}$) that effects state-level probabilities differently, we could specify this as,

$$
\begin{aligned}
\mathbf{z}_i &\sim \text{Categorical}(\boldsymbol{\psi_i}) \\
\boldsymbol{\psi}_i &= \begin{bmatrix} \psi_i^1 & \psi_i^2 & \psi_i^3 & \psi_i^4 \end{bmatrix}. \\
\psi_i^1 &= \frac{\phi_i^1}{\phi_i^1 + \phi_i^2 + \phi_i^3 + \phi_i^4} \\
\psi_i^2 &= \frac{\phi_i^2}{\phi_i^1 + \phi_i^2 + \phi_i^3 + \phi_i^4} \\
\psi_i^3 &= \frac{\phi_i^3}{\phi_i^1 + \phi_i^2 + \phi_i^3 + \phi_i^4} \\
\psi_i^4 &= \frac{\phi_i^4}{\phi_i^1 + \phi_i^2 + \phi_i^3 + \phi_i^4}.
\end{aligned}
$$

$$
\begin{aligned}
\phi_i^1 &= 1 \\
\phi_i^2 &= e^{\alpha_1 + \alpha_2 \times x_i} \\
\phi_i^3 &= e^{\alpha_3 + \alpha_4 \times x_i} \\
\phi_i^4 &= e^{\alpha_1 + \alpha_2 \times x_i + \alpha_3 + \alpha_4 \times x_i + \alpha_5 + \alpha_6 \times x_i}
\end{aligned}
$$

A diffuse prior could be $\alpha_\bullet \sim \text{Logistic}(0, 1)$. This model is coded for JAGS in the file 'jags.multistate.occ.full.site.covs.R'.

##### 1.1.4 Multinomial-logit Model 3

We could generalize the modeling of state-specific occupancy parameters using separate design matrices ($\mathbf{x}_i^{\text{Day}}, \mathbf{x}_i^{\text{Night}}, \mathbf{x}_i^{\text{ND}}$) that for each site $i$ are 1 x $\text{Q}_m$ (being the number of columns) and associated vectors of coefficients ($\boldsymbol{\alpha}^{\text{Day}}, \boldsymbol{\alpha}^{\text{Night}}, \boldsymbol{\alpha}^{\text{ND}}$) that are $Q_m$ x 1, and specified as,

$$
\begin{aligned}
\mathbf{z}_i &\sim \text{Categorical}(\boldsymbol{\psi_i}) \\
\boldsymbol{\psi}_i &= \begin{bmatrix} \psi_i^1 & \psi_i^2 & \psi_i^3 & \psi_i^4 \end{bmatrix}. \\
\psi_i^1 &= \frac{\phi_i^1}{\phi_i^1 + \phi_i^2 + \phi_i^3 + \phi_i^4} \\
\psi_i^2 &= \frac{\phi_i^2}{\phi_i^1 + \phi_i^2 + \phi_i^3 + \phi_i^4} \\
\psi_i^3 &= \frac{\phi_i^3}{\phi_i^1 + \phi_i^2 + \phi_i^3 + \phi_i^4} \\
\psi_i^4 &= \frac{\phi_i^4}{\phi_i^1 + \phi_i^2 + \phi_i^3 + \phi_i^4}.
\end{aligned}
$$

$$
\begin{aligned}
\phi_i^1 &= 1 \\
\phi_i^2 &= e^{\mathbf{x}_i^{\text{Day}}\boldsymbol{\alpha}^{\text{Day}}} \\
\phi_i^3 &= e^{\mathbf{x}_i^{\text{Night}}\boldsymbol{\alpha}^{\text{Night}}} \\
\phi_i^4 &= e^{\mathbf{x}_i^{\text{Day}}\boldsymbol{\alpha}^{\text{Day}}+\mathbf{x}_i^{\text{Night}}\boldsymbol{\alpha}^{\text{Night}}+\mathbf{x}_i^{\text{ND}}\boldsymbol{\alpha}^{\text{ND}}}
\end{aligned}
$$

A diffuse prior for all coefficients could be done using Logistic$(0, 1)$. This model allows state-specific modeling of occupancy probabilities by any combination of covaraites when specificed using a design matrix, which is common in linear regression modeling. This model is coded for JAGS in the file 'jags.multistate.occ.full.site.covs.R'.

##### 1.1.5 Multinomial-logit Model 2

We could also model detection parameters by site ($\mathbf{x}$) or survey specific ($\mathbf{V}$) covarariates, such that

$$
\begin{aligned}
\boldsymbol{q}_{ij} &= \begin{bmatrix} 1 & 0 & 0 & 0 \\ 1 & q_{ij}^{2,2} & 0 & 0 \\ 1 & 0 & q_{ij}^{3,3} & 0 \\ 1 & q_{ij}^{4,2} & q_{ij}^{4,3} & q_{ij}^{4,4} \end{bmatrix} \\
q_{ij}^{2,2} &= e^{\beta_0^{\text{Day}}+\beta_1^{\text{Day}}\times x_i+\beta_2^{\text{Day}}\times v_{ij}} \\
q_{ij}^{3,3} &= e^{\beta_0^{\text{Night}}+\beta_1^{\text{Night}}\times x_i+\beta_2^{\text{Day}}\times v_{ij}} \\
q_{ij}^{4,2} &= e^{\beta_0^{\text{ND2}}+\beta_1^{\text{ND2}}\times x_i+\beta_2^{\text{Day}}\times v_{ij}} \\
q_{ij}^{4,3} &= e^{\beta_0^{\text{ND3}}+\beta_1^{\text{ND3}}\times x_i+\beta_2^{\text{Day}}\times v_{ij}} \\
q_{ij}^{4,4} &= e^{\beta_0^{\text{ND4}}+\beta_1^{\text{ND4}}\times x_i+\beta_2^{\text{Day}}\times v_{ij}}
\end{aligned}
$$

and the probability of detection parameters can be derived as,

$$
\begin{aligned}
p_{ij}^{\text{Day}} &= \frac{q_{ij}^{2,2}}{1+q_{ij}^{2,2}+0+0} \\
p_{ij}^{\text{Night}} &= \frac{q_{ij}^{3,3}}{1+0+q_{ij}^{3,3}+0} \\
p_{ij}^{\text{ND1}} &= \frac{1}{1+q_{ij}^{4,2}+q_{ij}^{4,3}+q_{ij}^{4,4}} \\
p_{ij}^{\text{ND2}} &= \frac{q_{ij}^{4,2}}{1+q_{ij}^{4,2}+q_{ij}^{4,3}+q_{ij}^{4,4}} \\
p_{ij}^{\text{ND3}} &= \frac{q_{ij}^{4,3}}{1+q_{ij}^{4,2}+q_{ij}^{4,3}+q_{ij}^{4,4}} \\
p_{ij}^{\text{ND4}} &= \frac{q_{ij}^{4,4}}{1+q_{ij}^{4,2}+q_{ij}^{4,3}+q_{ij}^{4,4}}.
\end{aligned}
$$

Diffuse priors for all $\beta$ parameters could be specified as Logistic(0,1).

#### 1.2 Reduced MSDOM

##### 1.2.1 Probability Model

We might consider a simpler structure for $\boldsymbol{\psi}$ or $\boldsymbol{P}$ because of ecological or data sparsity reasons. It might be that sites used during the day and night (state 4) are not unique, but simply random and independently determined by the probability any site is used during the day ($\psi^{\text{Day.M}}$) and night ($\psi^{\text{Night.M}}$). Rather than exclusive state probabilities, $\psi^{\text{Day.M}}$ is the probability of using any site during the day, regardless of use or not use at night (marginal probability); further, $\psi^{\text{Night.M}}$ is the probability of using any site at night, regardless of use during the day. We can redefine our occupancy parameters as,

$$
\begin{aligned}
\mathbf{z}_i &\sim \text{Categorical}(\boldsymbol{\psi}) \\
\boldsymbol{\psi} &= \begin{bmatrix} \psi^1 & \psi^2 & \psi^3 & \psi^4 \end{bmatrix} \\
\psi^1 &= (1-\psi^{\text{Day.M}})(1-\psi^{\text{Night.M}}) \\
\psi^2 &= \psi^{\text{Day.M}}(1-\psi^{\text{Night.M}}) \\
\psi^3 &= (1-\psi^{\text{Day.M}})\psi^{\text{Night.M}} \\
\psi^4 &= \psi^{\text{Day.M}}\psi^{\text{Night.M}}.
\end{aligned}
$$

Similarly, we can simplify the detection matrix using the probability of detection during the day ($p^{\text{Day.M}}$) and night ($p^{\text{Night.M}}$) as,

$$
\boldsymbol{P} = \begin{bmatrix} 1 & 0 & 0 & 0 \\ 1-p^{\text{Day.M}} & p^{\text{Day.M}} & 0 & 0 \\ 1-p^{\text{Night.M}} & 0 & p^{\text{Night.M}} & 0 \\ (1-p^{\text{Day.M}})(1-p^{\text{Night.M}}) & p^{\text{Day.M}}(1-p^{\text{Night.M}}) & (1-p^{\text{Day.M}})p^{\text{Night.M}} & p^{\text{Day.M}}p^{\text{Night.M}} \end{bmatrix} \tag{5}
$$

, where $p^{1,4}$ can also be specified as equal to $1 - p^{\text{Day.M}}(1 - p^{\text{Night.M}}) - (1 - p^{\text{Day.M}})p^{\text{Night.M}} - p^{\text{Day.M}}p^{\text{Night.M}}$. Unless the species of interest is completely cathemeral and uses the day and night times in equal proportion, it is unlikely that $p^{\text{Day.M}}$ and $p^{\text{Night.M}}$ should be equivalent. We could use the diffuse prior distributions of

$$
\begin{aligned}
\psi^{\text{Day.M}} &\sim \text{Beta}(1,1) \\
\psi^{\text{Night.M}} &\sim \text{Beta}(1,1) \\
p^{\text{Day.M}} &\sim \text{Beta}(1,1) \\
p^{\text{Night.M}} &\sim \text{Beta}(1,1).
\end{aligned}
$$

This model is coded for JAGS in the file 'jags.multistate.occ.reduced.R'.

#### 1.3 Logit Model 1

We could consider the effects of a site-level covariate ($\mathbf{x}$) on the marginal occurrence probabilities ($\psi^{\text{Day.M}}, \psi^{\text{Night.M}}$) by specifying,

$$
\begin{aligned}
\mathbf{z}_i &\sim \text{Categorical}(\boldsymbol{\psi}_i) \\
\boldsymbol{\psi}_i &= \begin{bmatrix} \psi_i^1 & \psi_i^2 & \psi_i^3 & \psi_i^4 \end{bmatrix} \\
\psi_i^1 &= (1 - \psi_i^{\text{Day.M}})(1 - \psi_i^{\text{Night.M}}) \\
\psi_i^2 &= \psi_i^{\text{Day.M}}(1 - \psi_i^{\text{Night.M}}) \\
\psi_i^3 &= (1 - \psi_i^{\text{Day.M}})\psi_i^{\text{Night.M}} \\
\psi_i^4 &= \psi_i^{\text{Day.M}}\psi_i^{\text{Night.M}} \\
\text{logit}(\psi_i^{\text{Day.M}}) &= \alpha_1^{\text{Day}} + \alpha_2^{\text{Day}} \times x_i \\
\text{logit}(\psi_i^{\text{Night.M}}) &= \alpha_1^{\text{Night}} + \alpha_2^{\text{Night}} \times x_i
\end{aligned}
$$

Diffuse prior distributions for all $\alpha$ parameters could be Logistic$(0, 1)$. This model is coded for JAGS in the file 'jags.multistate.occ.reduced.site.covs.R'

##### 1.3.1 Logit Model 2

We could generalize the modeling of marginal occupancy parameters using separate design matrices ($\mathbf{x}_i^{\text{Day}}, \mathbf{x}_i^{\text{Night}}$) that for each site $i$ are 1 x $\text{Q}_m$ (being the number of columns) and associated vectors of coefficients ($\boldsymbol{\alpha}^{\text{Day}}, \boldsymbol{\alpha}^{\text{Night}}$) that are $Q_m$ x 1, and specified as,

$$
\begin{aligned}
\mathbf{z}_i &\sim \text{Categorical}(\boldsymbol{\psi_i}) \\
\boldsymbol{\psi}_i &= \begin{bmatrix} \psi_i^1 & \psi_i^2 & \psi_i^3 & \psi_i^4 \end{bmatrix}. \\
\psi_i^1 &= (1-\psi_i^{\text{Day.M}})(1-\psi_i^{\text{Night.M}}) \\
\psi_i^2 &= \psi_i^{\text{Day.M}}(1-\psi_i^{\text{Night.M}}) \\
\psi_i^3 &= (1-\psi_i^{\text{Day.M}})\psi_i^{\text{Night.M}} \\
\psi_i^4 &= \psi_i^{\text{Day.M}}\psi_i^{\text{Night.M}} \\
\text{logit}(\psi_i^{\text{Day.M}}) &= \mathbf{x}_i^{\text{Day}}\boldsymbol{\alpha}^{\text{Day}} \\
\text{logit}(\psi_i^{\text{Night.M}}) &= \mathbf{x}_i^{\text{Night}}\boldsymbol{\alpha}^{\text{Night}}
\end{aligned}
$$

Diffuse prior distributions for all $\alpha$ parameters could be Logistic$(0, 1)$. This model is coded for JAGS in the file 'jags.multistate.occ.reduced.site.covs.by.state.R'.

##### 1.3.2 Multinomial-logit Mode 1

An alternative reduced model specification is to use the multinomial-logit parameterization, which can constrain relationships among state probabilities and include covariates. This model could be parameterized as,

$$
\begin{aligned}
\mathbf{z}_i &\sim \text{Categorical}(\boldsymbol{\psi_i}) \\
\boldsymbol{\psi}_i &= \begin{bmatrix} \psi_i^1 & \psi_i^2 & \psi_i^3 & \psi_i^4 \end{bmatrix}. \\
\psi_i^1 &= \frac{\phi_i^1}{\phi_i^1+\phi_i^2+\phi_i^3+\phi_i^4} \\
\psi_i^2 &= \frac{\phi_i^2}{\phi_i^1+\phi_i^2+\phi_i^3+\phi_i^4} \\
\psi_i^3 &= \frac{\phi_i^3}{\phi_i^1+\phi_i^2+\phi_i^3+\phi_i^4} \\
\psi_i^4 &= \frac{\phi_i^4}{\phi_i^1+\phi_i^2+\phi_i^3+\phi_i^4}. \\
\phi_i^1 &= 1 \\
\phi_i^2 &= e^{\alpha_1} \\
\phi_i^3 &= e^{\alpha_2} \\
\phi_i^4 &= e^{\alpha_1+\alpha_2}
\end{aligned}
$$

The detection matrix could also be specified similarly using the multinomial-logit as,

$$
\boldsymbol{q}_j = \begin{bmatrix} 1 & 0 & 0 & 0 \\ 1 & e^{\beta_1} & 0 & 0 \\ 1 & 0 & e^{\beta_2} & 0 \\ 1 & e^{\beta_1} & e^{\beta_2} & e^{\beta_1+\beta_2} \end{bmatrix} \tag{6}
$$

and where the detection parameters are derived as,

$$
\begin{aligned}
p^{\text{Day}} &= \frac{e^{\beta_1}}{1 + e^{\beta_1} + 0 + 0} \\
p^{\text{Night}} &= \frac{e^{\beta_2}}{1 + 0 + e^{\beta_2} + 0} \\
p^{\text{ND4}} &= \frac{e^{\beta_1+\beta 2}}{1 + e^{\beta_1} + e^{\beta_2} + e^{\beta_1+\beta 2}}
\end{aligned}
$$

This model could be easily extended to include a difference of occupancy and detection in state 4 by a small modification of $\phi_i^4 = e^{\alpha_1+\alpha_2+\alpha_3}$ and the matrix cell $q^{4,4}$ to $e^{\beta_1+\beta_2+\beta_3}$. This would constrain the probability of detection during the day to be the same in state 2 and state 4 and the probabilty of detection during the night to be the same in state 3 and state 4. But, detection of state 4 in state 4 would be allowed to be more or less than the combination of the probability of detection during the day and night alone. A diffuse prior for all coefficients could be done using Logistic$(0, 1)$.

#### 1.4 Null MSDOM

##### 1.4.1 Probability Model

Furthermore, we could consider a multi-state model in which there is no spatial variability in temporal occurrence. This could be used as a null model in a model comparison framework to assess the importance of this type of heterogeneity. The state-specific occurrence probabilities can be described by the overall or marginal (regardless of state) occurrence probability ($\psi^\bullet$); since we assume the states do not matter, we then assume the probability of occurrence in each state is the same. The same is true for the overall or marginal probability of detection ($p^\bullet$). This model would be essentially a single state (occurrence or not occurrence) model. We can do so by defining,

$$
\begin{aligned}
\mathbf{z}_i &\sim \text{Categorical}(\boldsymbol{\psi}) \\
\boldsymbol{\psi} &= \begin{bmatrix}\psi^1 & \psi^2 & \psi^3 & \psi^4\end{bmatrix}. \\
\psi^1 &= 1 - \psi^{\bullet} \\
\psi^2 &= \frac{\psi^{\bullet}}{3} \\
\psi^3 &= \frac{\psi^{\bullet}}{3} \\
\psi^4 &= \frac{\psi^{\bullet}}{3} \\
\boldsymbol{\psi} &= \begin{bmatrix}\psi^1 & \psi^2 & \psi^3 & \psi^4\end{bmatrix}.
\end{aligned}
$$

Similarly, we can define the detection matrix using $p^{\bullet}$ as,

$$
\boldsymbol{p}_j = \begin{bmatrix} 1 & 0 & 0 & 0 \\ 1 - p_j^{\bullet} & p_j^{\bullet} & 0 & 0 \\ 1 - p_j^{\bullet} & 0 & p_j^{\bullet} & 0 \\ 1 - p_j^{\bullet} & \frac{p_j^{\bullet}}{3} & \frac{p_j^{\bullet}}{3} & \frac{p_j^{\bullet}}{3} \end{bmatrix} \tag{7}
$$

We could use the diffuse prior distributions of

$$
\begin{aligned}
\psi^{\bullet} &\sim \text{Beta}(1, 1) \\
p_j^{\bullet} &\sim \text{Beta}(1, 1).
\end{aligned}
$$

This model is coded for JAGS in the file ‘jags.multistate.occ.null.R’.

##### 1.4.2 Multinomial-logit Model 1

An alternative specification uses the multinomial logit to transform occurrence and detection probabilities to the logit-scale while ensuring appropriate constrains on parameters (i.e., probabilities stay between 0 and 1). This model specification is more conducive to including covariates to model their effects on different parameters. To do so we can define occupancy parameters with a shared log-valued parameter ($\alpha$), such that

$$
\begin{aligned}
\mathbf{z}_i &\sim \text{Categorical}(\boldsymbol{\psi}) \\
\boldsymbol{\psi} &= \begin{bmatrix}\psi^1 & \psi^2 & \psi^3 & \psi^4\end{bmatrix} \\
\psi^1 &= \frac{\phi^1}{\phi^1+\phi^2+\phi^3+\phi^4} \\
\psi^2 &= \frac{\phi^2}{\phi^1+\phi^2+\phi^3+\phi^4} \\
\psi^3 &= \frac{\phi^3}{\phi^1+\phi^2+\phi^3+\phi^4} \\
\psi^4 &= \frac{\phi^4}{\phi^1+\phi^2+\phi^3+\phi^4}. \\
\phi^1 &= 3 \\
\phi^2 &= e^\alpha \\
\phi^3 &= e^\alpha \\
\phi^4 &= e^\alpha
\end{aligned}
$$

Similarly, we can define the detection matrix on the logit scale using a shared log-valued parameter ($\beta$), such that

$$
\boldsymbol{q}_j = \begin{bmatrix} 1 & 0 & 0 & 0 \\ 1 & e^\beta & 0 & 0 \\ 1 & 0 & e^\beta & 0 \\ 3 & e^\beta & e^\beta & e^\beta \end{bmatrix}. \tag{8}
$$

The probability of detection can be derived as,

$$
p^\bullet = \frac{e^\beta}{1+e^\beta+0+0} \tag{9}
$$

We could use the diffuse prior distributions of

$$
\begin{aligned}
\alpha &\sim \text{Logistic}(0,1) \\
\beta &\sim \text{Logistic}(0,1).
\end{aligned}
$$

##### 1.4.3 Multinomial-logit Model 2

We could consider the effects of a site-level covariate ($\mathbf{x}$) on the occupancy probability by specifying,

$$
\begin{aligned}
\mathbf{z}_i &\sim \text{Categorical}(\boldsymbol{\psi}_i) \\
\boldsymbol{\psi}_i &= \begin{bmatrix} \psi_i^1 & \psi_i^2 & \psi_i^3 & \psi_i^4 \end{bmatrix} \\
\psi_i^1 &= \frac{\phi^1}{\phi^1 + \phi^2 + \phi^3 + \phi^4} \\
\psi_i^2 &= \frac{\phi^2}{\phi^1 + \phi^2 + \phi^3 + \phi^4} \\
\psi_i^3 &= \frac{\phi^3}{\phi^1 + \phi^2 + \phi^3 + \phi^4} \\
\psi_i^4 &= \frac{\phi^4}{\phi^1 + \phi^2 + \phi^3 + \phi^4}. \\
\phi_i^1 &= 3 \\
\phi_i^2 &= e^{\alpha_1 + \alpha_2 \times x_i} \\
\phi_i^3 &= e^{\alpha_1 + \alpha_2 \times x_i} \\
\phi_i^4 &= e^{\alpha_1 + \alpha_2 \times x_i}
\end{aligned}
$$

Since we are ignoring the state designations, there is no reason to use different design matrices of each state-occupancy parameter nor a reason to estimate different parameters. This model is coded for JAGS in the file 'jags.multistate.occ.null.site.covs.R'.

##### 1.4.4 Multinomial-logit Model 3

We could generalize the modeling of the occupancy parameter using the same design matrix ($\mathbf{x}_i$) that for each site $i$ that is 1 x Q (being the number of columns) and associated vectors of coefficients ($\boldsymbol{\alpha}$) that are $Q_m$ x 1, we specify

$$
\begin{aligned}
\mathbf{z}_i &\sim \text{Categorical}(\boldsymbol{\psi}_i)\\
\boldsymbol{\psi}_i &= \begin{bmatrix}\psi_i^1 & \psi_i^2 & \psi_i^3 & \psi_i^4\end{bmatrix}\\
\psi_i^1 &= \frac{\phi^1}{\phi^1+\phi^2+\phi^3+\phi^4}\\
\psi_i^2 &= \frac{\phi^2}{\phi^1+\phi^2+\phi^3+\phi^4}\\
\psi_i^3 &= \frac{\phi^3}{\phi^1+\phi^2+\phi^3+\phi^4}\\
\psi_i^4 &= \frac{\phi^4}{\phi^1+\phi^2+\phi^3+\phi^4}.\\
\phi_i^1 &= 3\\
\phi_i^2 &= e^{\mathbf{x}_i\boldsymbol{\alpha}}\\
\phi_i^3 &= e^{\mathbf{x}_i\boldsymbol{\alpha}}\\
\phi_i^4 &= e^{\mathbf{x}_i\boldsymbol{\alpha}}.
\end{aligned}
$$

Since we are ignoring the state designations, there is no reason to use different design matrices of each state-occupancy parameter nor a reason to estimate different parameters. This model is coded for JAGS in the file 'jags.multistate.occ.null.site.covs.by.state.R'.

### 2 Model Selection

To compare models (e.g., Full, Reduced, and Null MSDOM) we could use the conditional predictive ordinate (CPO; Hooten and Hobbs 2015). To do so, we need to define the integrated likelihood. First, lets define $I_i$ as an indicator for site $i$ that is a 1 when the logical argument is true and otherwise 0 (see below) and $\boldsymbol{f}_i = \begin{bmatrix} f_i^1 f_i^2 f_i^3 f_i^4 \end{bmatrix}$ is matrix of the frequency of observations in each state across all $K$ surveys. Therefore, assuming detection parameters are constant over each $j$ survey, the integrated likelihood is specified as,

$$
\begin{aligned}
[\mathbf{y}_i|\boldsymbol{\psi}_i,\boldsymbol{p}] = &\left(\psi^1+\psi^2(1-p^{2,2})^{f_i^1}+\psi^3(1-p^{3,3})^{f_i^1}+\psi^4(1-p^{1,4})^{f_i^1}\right)^{I_{\max(\boldsymbol{y}_{i\cdot})\equiv 1}}\\
&\times\left(\psi^2(p^{2,2})^{f_i^2}(1-p^{2,2})^{f_i^1}+\psi^4(p^{1,4})^{f_i^1}(p^{2,4})^{f_i^2}(p^{3,4})^{f_i^3}(p^{4,4})^{f_i^4}\right)^{I_{\max(\boldsymbol{y}_{i\cdot})\equiv 2}}\\
&\times\left(\psi^3(p^{3,3})^{f_i^3}(1-p^{3,3})^{f_i^1}+\psi^4(p^{1,4})^{f_i^1}(p^{2,4})^{f_i^2}(p^{3,4})^{f_i^3}(p^{4,4})^{f_i^4}\right)^{I_{\max(\boldsymbol{y}_{i\cdot})\equiv 3\wedge\forall\boldsymbol{y}_{i\cdot}\not\equiv 2}}\\
&\times\left(\psi^4(p^{1,4})^{f_i^1}(p^{2,4})^{f_i^2}(p^{3,4})^{f_i^3}(p^{4,4})^{f_i^4}\right)^{I_{\max(\boldsymbol{y}_{i\cdot})\equiv 4\vee 2\in\boldsymbol{y}_{i\cdot}\wedge 3\in\boldsymbol{y}_{i\cdot}}}
\end{aligned}
$$

Following **?**, for each MCMC iteration $s = 1,\ldots, S$,

$$\text{CPO} = -\sum_{i=1}^{N} \log(CPO_{ij})$$

$$\text{CPO}_{ij} = \frac{S}{\sum_{s=1}^{S} [\mathbf{y}_i | \boldsymbol{\psi}_i^{(s)}, \boldsymbol{p}^{(s)}]^{-1}}$$

For models with or without site-level variation in state occurrence and no site or survey level variation in detection, CPO can be calculated for these models using the function 'CPO.function.r'.

### 3 Goodness of fit

We can evaluate the overall fit of the model using a Bayesian p-value comparing the observed ($D$) and predicted model deviance ($\tilde{D}$) as,

$$D^{(s)} = -2 \cdot \sum_{i=1}^{N} \log[\mathbf{y}_i | \boldsymbol{\psi}_i^{(s)}, \boldsymbol{p}^{(s)}]$$

$$\tilde{D}^{(s)} = -2 \cdot \sum_{i=1}^{N} \log[\tilde{\mathbf{y}}_i | \boldsymbol{\psi}_i^{(s)}, \boldsymbol{p}^{(s)}]$$

where the predicted occurrences and detections are sampled for each $s$ MCMC iteration as,

$$\tilde{\boldsymbol{z}}_i^{(s)} \sim \text{Categorical}(\boldsymbol{\psi}_i^{(s)})$$

$$\tilde{\boldsymbol{y}}_{ij}^{(s)} \sim \text{Categorical}(\tilde{\mathbf{z}}_i \cdot \boldsymbol{p}_j)$$

The Bayesian P-value can be calculated as the proportion of times $D$ is greater than $\tilde{D}$. For a model that fits well, the p-value should be near 0.5; if the p-value is greater than 0.95 or less than 0.05 than the model foes not fit the data well.
