## Appendix S3 for "Rethinking habitat occupancy modeling and the role of diel activity in an anthropogenic world" (from another version of this preprint: version 1, 2021-07-01)

### Appendix S3: Additional data collection details and methodology

### 1. Fosa Dataset

| Region | Site ID | Year | Total Cameras | Survey Dates | Season |
|---|---|---|---|---|---|
| Makira | SITE 1 | 2008 | 20 | Sept 2 – Nov 13 | Cool, Dry |
| | SITE 1 | 2010 | 24 | Sept 16 – Nov 16 | Cool, Dry |
| | SITE 1 | 2011 | 24 | Aug 20 – Oct 20 | Cool, Dry |
| | SITE 1 | 2012 | 24 | Aug 1 – Oct 16 | Cool, Dry |
| | SITE 1 | 2013 | 24 | Sept 7 – Nov 20 | Cool, Dry |
| | SITE 1 | 2015 | 24 | Sept 11 – Nov 9 | Cool, Dry |
| | SITE 2 | 2011 | 24 | Mar 20 – May 23 | Warm, Wet |
| | SITE 2 | 2012 | 24 | Jun 9 – July 23 | Cold, Dry |
| | SITE 2 | 2013-2014 | 24 | Nov 16 – Jan 7 | Hot, Wet |
| | SITE 3 | 2009 | 20 | Mar 4 – May 4 | Warm, Wet |
| | SITE 4 | 2009 | 19 | Aug 21 – Oct 27 | Cool, Dry |
| | SITE 5 | 2009-2010 | 18 | Nov 20 – Jan 27 | Hot, Wet |
| | SITE 6 | 2010-2011 | 24 | Dec 9 – Feb 18 | Hot, Wet |
| | SITE 7 | 2011 | 24 | Jun 9 – Aug 13 | Cold, Dry |
| Ranomafana | VAL-SAH | 2008 | 53 | June – Oct | Cold, Dry |
| | SAH-CVB | 2007 | 42 | June – Aug | Cold, Dry |

Table 1. Details on seven survey regions used for analysis of fosa in the Makira Natural Park and Ranomafana National Park Regions. Survey dates refer to the first date cameras were set for the specific site and year followed by the last date cameras were removed.

**Incorporating multiple camera surveys for the fosa case studies**

We modeled the variation among data collection years and survey areas differently for the Makira and Ranomafana datasets. For the Makira data, we accounted for this variation by treating each model parameter as a random effect across survey areas (e.g., for s survey areas, $\alpha_s^{Day} \sim \text{Normal}(\mu_{\alpha^{Day}}, \sigma_{\alpha^{Day}})$ ); as such, model parameter estimates reported in the main text are the population-level mean effects (inference for across survey areas for each parameter). We did not treat each survey independently as this would allow 'Site 1' to have an oversized influence on parameter estimates as it was surveyed six times. Survey areas with multiple years of surveys (e.g., 'Site 1') were modeled together based on preliminary findings for a lack of difference among years. Specifically, we evaluated across year differences within a survey area by treating surveys as a categorical variable in a stacked occupancy design (see Monterroso et al., 2020). Models without survey year variation were more supported by CPO. With only two surveys at Ranomafana, we only used a stacked occupancy design, treating survey as a categorical variable, and compare models with and without this variable.

| Model | Model description | CPO | Δ CPO |
|---|---|---|---|
| *M1.full* | Full model with distance to town covariate | 838.50 | 0.00 |
| *M4.red* | Reduced model with distance to matrix covariate | 843.12 | 4.62 |
| *M9.red.no.covs* | Reduced model with no covariates | 843.64 | 5.14 |
| *M8.full2.no.covs* | Full model with no covariates and categorical effect | 843.74 | 5.24 |
| *M2.full* | Full model with distance to matrix covariate | 843.76 | 5.26 |
| *M4.red2* | Reduced model with distance to matrix covariate and categorical effect | 844.58 | 6.08 |
| *M3.red* | Reduced model with distance to town covariate | 844.65 | 6.15 |
| *M2.full2* | Full model with distance to matrix covariate and categorical effect | 844.68 | 6.18 |
| *M9.red2.no.covs* | Reduced model with no covariates and categorical effect | 844.82 | 6.32 |
| *M1.full2* | Full model with distance to town covariate and categorical effect | 845.50 | 7.00 |
| *M3.red2* | Reduced model with distance to town covariate and categorical effect | 845.99 | 7.49 |
| *M8.full.no.covs* | Full model with no covariates | 846.11 | 7.61 |
| *M6.null* | Null model with distance to matrix covariate | 916.25 | 77.75 |
| *M7.null* | Null model with no covariates | 916.27 | 77.77 |
| *M5.null* | Null model with distance to town covariate | 917.00 | 78.50 |
| *M7.null2* | Null model with no covariates and categorical effect | 917.00 | 78.50 |
| *M6.null2* | Null model with distance to matrix covariate and categorical effect | 917.19 | 78.69 |
| *M5.null2* | Null model with distance to town covariate and categorical effect | 917.80 | 79.295612 |

Table 2. Conditional predictive ordinate outputs for Ranomafana data. Lower values indicate a more supported model.

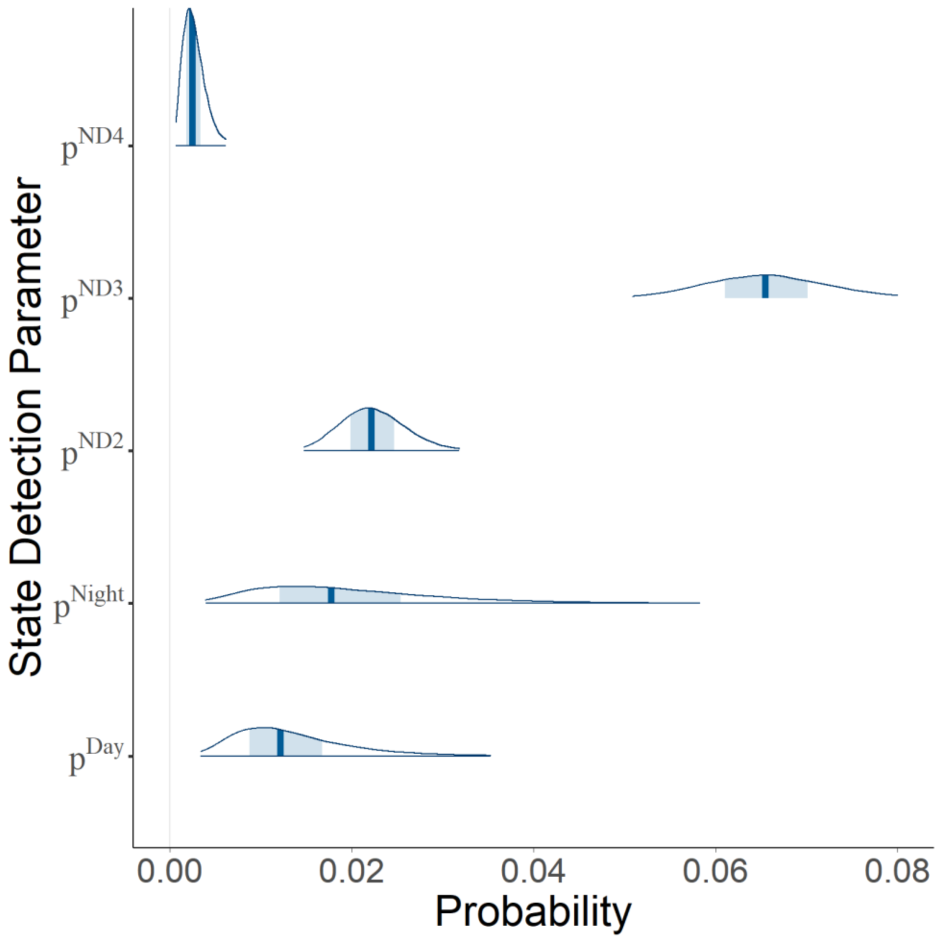


Figure 1. Posterior distributions of fosa state detection probability for the most supported model using the Ranomafana National Park data. The light blue shaded area represents 50% probability density and the dark blue line indicates the posterior mode. Note, $p^{ND1}$ is not plotted.

| | | Credible Interval | |
|---|---|---|---|
| **Parameter** | **Mode** | **Lower 95%** | **Upper 95%** |
| $\boldsymbol{p}^{\mathbf{Day}}$ | 0.01 | 0.01 | 0.00 |
| $\boldsymbol{p}^{\mathbf{Night}}$ | 0.02 | 0.02 | 0.00 |
| $\boldsymbol{p}^{\mathbf{ND1}}$ | 0.91 | 0.91 | 0.89 |
| $\boldsymbol{p}^{\boldsymbol{ND}\mathbf{2}}$ | 0.02 | 0.02 | 0.02 |
| $\boldsymbol{p}^{\boldsymbol{ND}\mathbf{3}}$ | 0.07 | 0.07 | 0.05 |
| $\boldsymbol{p}^{\boldsymbol{ND}\mathbf{4}}$ | 0.00 | 0.00 | 0.00 |

Table 3. Posterior quantities for detection probabilities of Ranomafana National Park from the best fit model. Credible intervals were calculated using the highest posterior density interval.

| **Model** | **Model description** | **CPO** | **Δ CPO** |
|---|---|---|---|
| *M1.full.no.covs* | Full model with no covariates | 1155.65 | 0.00 |
| *M2.full.covs* | Full model with covariates | 1158.05 | 2.40 |
| *M1.red.no.covs* | Reduced model with no covariates | 1173.68 | 18.03 |
| *M2.red.covs* | Reduced model with covariates | 1175.35 | 19.70 |
| *M1.null.no.covs* | Null model with covariates | 1215.40 | 59.75 |
| *M2.null.covs* | Null model with no covariates | 1216.78 | 61.13 |

Table 4. Conditional predictive ordinate outputs for Makira data. Lower values indicate a more supported model.

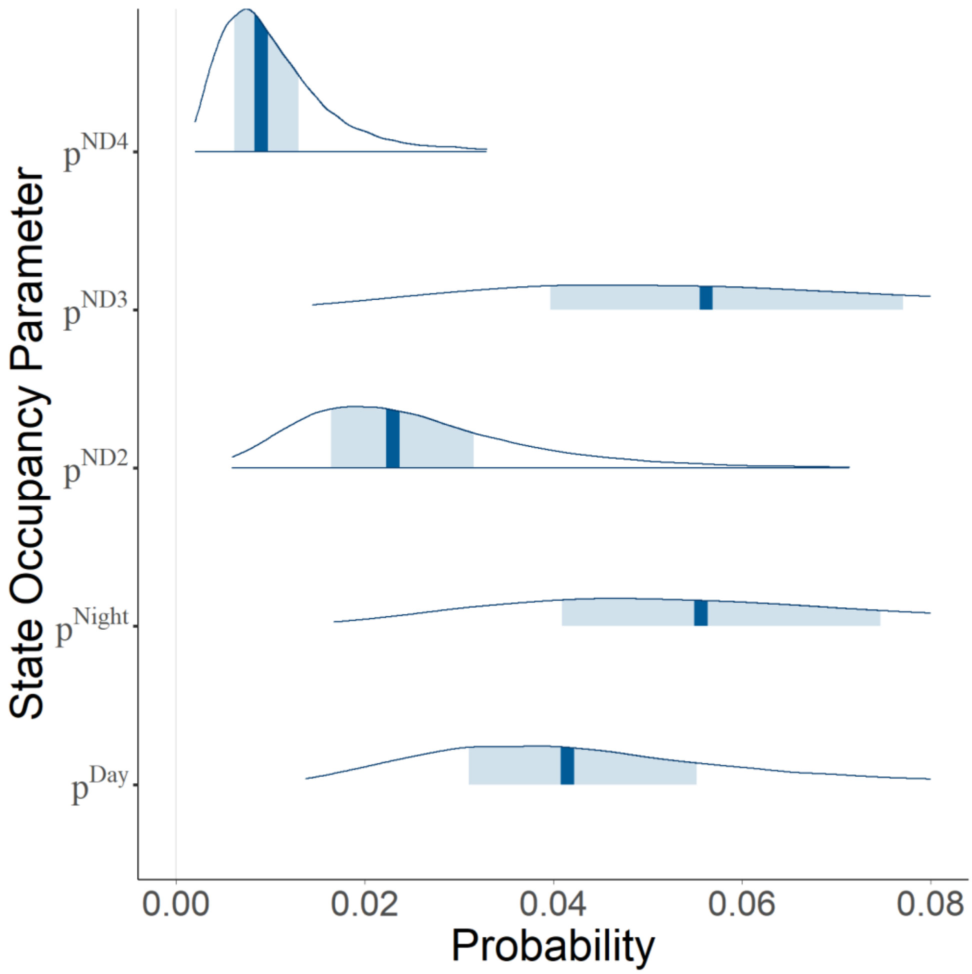


Figure 2. Posterior distributions of fosa state detection probability for the most supported model using the Makira Natural Park data. The light blue shaded area represents 50% probability density and the dark blue line indicates the posterior mode. Note, $p^{ND1}$ is not plotted.

| | | Credible Interval | |
|---|---|---|---|
| **Parameter** | **Mode** | **Lower 95%** | **Upper 95%** |
| $\boldsymbol{p}^{\mathbf{Day}}$ | 0.04 | 0.01 | 0.09 |
| $\boldsymbol{p}^{\mathbf{Night}}$ | 0.05 | 0.02 | 0.12 |
| $\boldsymbol{p}^{\mathbf{ND1}}$ | 0.29 | 0.14 | 0.47 |
| $\boldsymbol{p}^{\boldsymbol{ND}\mathbf{2}}$ | 0.02 | 0 | 0.05 |
| $\boldsymbol{p}^{\boldsymbol{ND}\mathbf{3}}$ | 0.05 | 0.01 | 0.12 |
| $\boldsymbol{p}^{\boldsymbol{ND}\mathbf{4}}$ | 0.01 | 0 | 0.02 |

Table 5. Posterior results for detection probabilities of the Makira region from the best fit model. Credible intervals were calculated using the highest posterior density interval.

### 2. Coyote Dataset

| | |
|---|---|
| **Camera** | Bushnell Trophy Cam Standard Edition and HD models |
| **Model numbers (and their count)** | 1199435 (8), 119436 (12), 1193537C (2), 119363C (19) |
| **Mode** | Camera |
| **Image Size:** | 5M Pixel |
| **Capture Number** | 1 |
| **Photo Video Size** | NA |
| **Video Length** | NA |
| **Interval** | 30 seconds |
| **Sensor Level** | Normal |
| **Format** | Execute (format memory card every time before deploying camera trap or replacing memory card) |
| **TV Out** | NTSC |
| **Time Stamp** | On |
| **Set Clock** | 24hr, year-month-day |
| **Field Scan** | Off |
| **Video Sound** | NA |
| **Default Set** | Cancel |

Table 6. Settings of motion-triggered camera traps and examples of lure treatments used in
The camera trap settings used on the camera traps for Fidino et al., The Effect of Lure on Detecting

Mammals with Camera Traps. Wildlife Society Bulletin.

**Data collection for Dynamic Modeling**

To capture seasonal variation in occupancy, cameras were placed for a minimum of 4 consecutive weeks in January, April, July, and October. Cameras were placed in urban greenspace such as city parks, cemeteries, golf courses, and natural areas, and sampling locations were 1 km apart at a minimum.
